# Designing Novel Solenoid Proteins with In Silico Evolution

**DOI:** 10.1101/2025.04.23.646631

**Authors:** Daniella Pretorius, Georgi I. Nikov, Kono Washio, Steve-William Florent, Henry N. Taunt, Sergey Ovchinnikov, James W. Murray

## Abstract

Solenoid proteins are elongated tandem repeat proteins with diverse biological functions, making them attractive targets for protein design. Advances in machine learning have transformed our understanding of sequence-structure relationships, enabling new approaches for de novo protein design. Here, we present an in silico evolution platform that couples a solenoid discriminator network with AlphaFold2 as an oracle within a genetic algorithm. Starting from random sequences, we design α-, β-, and αβ-solenoid backbones, generating structures that span natural and novel solenoid space. We experimentally characterise 41 solenoid designs, with α-solenoids consistently folding as intended, including one structurally validated design that closely matches the design model. All β-solenoids initially failed, reflecting the difficulty of designing β-strand majority proteins. By introducing terminal capping elements and refining designs based on earlier experimental screens, we generate two β-solenoids that have biophysical properties consistent with their designs. Our approach achieves fold-specific hallucination-based design without depending on explicit structural templates.

## Introduction

Solenoid proteins, a type of tandem repeat protein, are abundant in Nature. Tandem repeat proteins comprise about 14% of all proteins [1], with solenoid proteins specifically making up around 3% of the 56 million eukaryotic proteins in UniProt with AlphaFold2 (AF2)-predicted models [2]. These proteins perform diverse biological roles, driven by their modular helical shapes and secondary structure composition, which include α-, β-, and αβ-solenoids. For example, natural solenoid proteins function as antifreeze proteins [3] and DNA-binding proteins [4].

Solenoid proteins are attractive targets for de novo design due to their modular repeat architecture, which reduces sequence and structure complexity space. Their extended surfaces and helicity make them versatile for applications like peptide-binding [5], nano materials [6] and polyhedral nanocages [7]. However, traditional methods for designing new solenoid proteins have been slow and labor-intensive [8-10]. Recent advances in protein structure prediction with AF2 [11], RoseTTAFold [12] and trRosetta [13] have given rise to a new generation of protein design methods that repurpose these models to generate novel proteins, including diffusion and hallucination approaches.

Diffusion-based methods, such as RFdiffusion [14], add noise to protein structures and learn to reverse this process, generating plausible protein backbones from random Gaussian noise. However, their success often depends on post-design filtering with AF2, with many design trajectories being discarded. Hallucination methods repurpose structure prediction networks to iteratively optimise protein sequences and backbones [15-20]. Hallucination directly optimises sequences with structure prediction at each iteration, whereas diffusion-based methods generate samples from a distribution and rely on structure prediction for post-design filtering. Recent applications of hallucination have extended beyond unconditional protein generation to create symmetric oligomers [17], protein binders [19, 20], and functional motif scaffolding [16, 18]. While hallucination has also been applied to produce αβ repeat proteins by constraining secondary structure content [21], or refining luciferases starting from natural sequences [18], hallucination has yet to be applied to generating specific folds de novo.

Specifying a fold during design can be valuable, as certain fold classes have desirable properties that can be modified or integrated into new scaffolds. For example, de novo enzyme design often involves exploring specific fold spaces to generate proteins that perform natural catalytic functions with enhanced efficiency, modified specificity, or improved stability [18]. Diffusion-based approaches can condition on fold information explicitly using structural priors such as secondary structure and a coarse-grained description of the fold topology [14, 22, 23].

Here, we present an in silico protein evolution platform for solenoid protein design. The platform is guided by a solenoid discrimination network [24] in a hallucination loop with AF2 as an oracle, where sequences are updated using a genetic algorithm. Unlike other approaches that use explicit structural priors, this approach drives solenoid fold specificity using the discrimination network, enabling exploration of diverse and novel solenoid architectures. Using this platform, we generate a library of α-, β-, and αβ-solenoid proteins spanning both natural and novel structures. Five designs were highly stable, having biophysical properties consistent with their intended structures, yielding a 20% experimental success rate for the in silico evolution platform. One α-solenoid design was structurally characterised with X-ray crystallography. While α-solenoid designs were successful, all the initial β-solenoid designs aggregated or did not express. By introducing terminal capping regions to the initial designs, we generated two β-solenoid proteins that have biophysical properties consistent with their designed structures. These results highlight the difficulty of designing β-strand majority proteins, which to date have relied heavily on rules-based approaches [25-27].

## Results

### In silico evolution platform for solenoid protein design

We developed an in silico evolution platform for fold-specific protein generation, starting from random repeat sequences (**Fig. 1, Supplementary Fig. 1**). The platform is guided by a solenoid discrimination network, SOLeNNoID [24], which assigns per-residue solenoid class scores. We used a hallucination loop with AF2 as an oracle, where sequences are updated using a genetic algorithm. Using this approach, we designed α-, β-, and αβ-solenoid proteins, optimising for both AF2 pLDDT and solenoid score. In the in silico evolution platform, the sequence is a concatamer of N copies of a repeat of length L (**Fig. 1a**).

**Figure 1.**
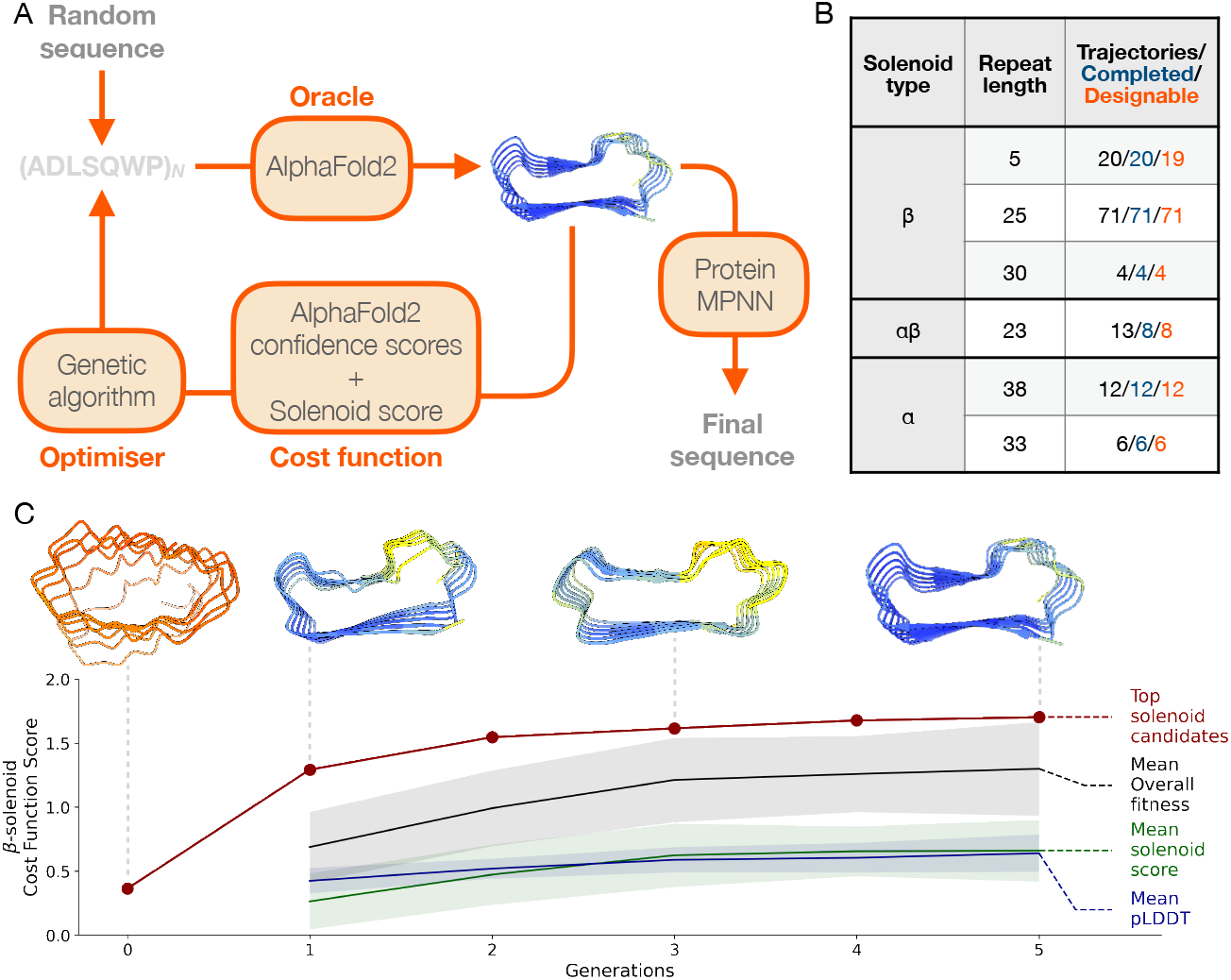
In silico evolution platform for solenoid protein generation. **(A)** The input repeat sequence of length L is repeated N times to make the full NxL sequence, whose structure is then predicted with AF2. The resulting design is scored with a cost function that combines AF2 pLDDT, as well as a solenoid-type probability score. The sequence pool is updated using a genetic algorithm. This continues until a set threshold for the cost function is met, and the output protein structure is taken forward. The sequence is redesigned using ProteinMPNN for successful backbones obtained through the in silico evolution platform. **(B)** Outcomes of different solenoid category and repeat lengths for solenoids generated with the *in silico* evolution platform **(C)** Example of trajectory for β-solenoid from an initial random sequence to final solenoid design. Generation 0 is the initial random starting sequence. The cost function value of the best candidate (*red*) is shown for each generation. The mean fitness (*black*), pLDDT (*blue*) and solenoid score (*green*) for each generation is shown. 1 standard deviation around the mean is shown by the shaded regions. Top candidate structures colored by pLDDT score are shown for generations 0, 1, 3, 5.

The cost function guiding the solenoid design was a weighted sum of AF2 pLDDT and SOLeNNoID scores, optimising for a selected solenoid type (α, β, or αβ) (**Fig. 1a, b**). Both scores range from 0 to 1, with a minimum pLDDT threshold of 0.7 applied to ensure high-confident AF2 predictions [11]. For α- and β-solenoids, a solenoid score threshold of 0.6 was used, while αβ-solenoids required a stricter threshold of 0.7. Lower thresholds frequently resulted in αβ-solenoid structures that had excessive disordered loops and resembled β-solenoids.

The efficiency of backbone generation varied across solenoid types and repeat lengths (**Fig. 1b**). Solenoids with smaller repeat lengths tended to converge on a high confidence solenoid structure faster, likely due to the reduced sequence space. For example, the platform generated ∼6 times more 25mer β-solenoids compared to 38mer α-solenoids, over a similar computational runtime. β-solenoids demonstrated rapid convergence, with some trajectories producing structures that satisfy the threshold criteria in as few as five generations (**Fig. 1c**).

Sequences generated during in silico evolution were redesigned with ProteinMPNN over the full NxL sequence length (**Fig. 1a**), addressing two key challenges. First, sequence redesign after hallucination has been shown to greatly improve experimental outcomes in previous work [16, 17]. The sequences generated via a hallucination method often have unrealistic properties. Sequences may also be adversarial, in that they “fool” the structure prediction model into producing confidently incorrect predictions [28]. For example, the sequences generated prior to redesign in this work showed a high abundance of cysteine residues, surface-exposed hydrophobic amino acids, and buried hydrophilic amino acids. Second, the hallucinated backbones contained identical repeats, which are often unsuitable for gene synthesis. Sequence redesign introduced variation, making them more realistic and synthesis-compatible.

Each redesigned sequence was then repredicted with AF2, and the resulting structures were ranked based on their pLDDT and RMSD relative to the original backbone. Design trajectories were tracked to evaluate the number initiated, that met the completion criteria within 30 generations, and resulted in backbones classified as “designable” (**Fig. 1b**). Designability was assessed by evaluating the consistency between the redesigned sequences and the original backbones. A backbone was classified as designable if any redesigned sequence yielded an AF2-predicted structure within 5 Å RMSD of the original backbone and an average pLDDT greater than 80. Using the platform, we generated 18 α-, 94 β-, and 8 αβ-solenoid designable backbones, achieving an overall designability rate of 99% (**Fig. 1b**). This exceeds the less than 1% designability rate reported for RFdiffusion when fold conditioning is applied [14].

### Designed solenoids span natural and novel solenoid space

A key question was whether the in silico evolution platform would generate novel solenoid structures or only recapitulate known ones. To assess this, we examined solenoid structure space using Multidimensional Scaling (MDS), based on pairwise structural alignments between our designed solenoids and solenoids in the PDB (**Fig. 2a**). In this representation, proximity between points corresponds to structural similarity. The MDS plot shows that our designed solenoids populate both known and unobserved regions of solenoid structural space, indicating that the platform generalises beyond natural solenoids in the PDB (**Fig. 2a**). Additionally, it was unclear whether identical repeat lengths would yield similar structures. However, substantial diversity was observed even within designs of the same repeat length. The MDS representation of solenoid space shows a clear separation between different solenoid types, whether natural or designed.

**Figure 2.**
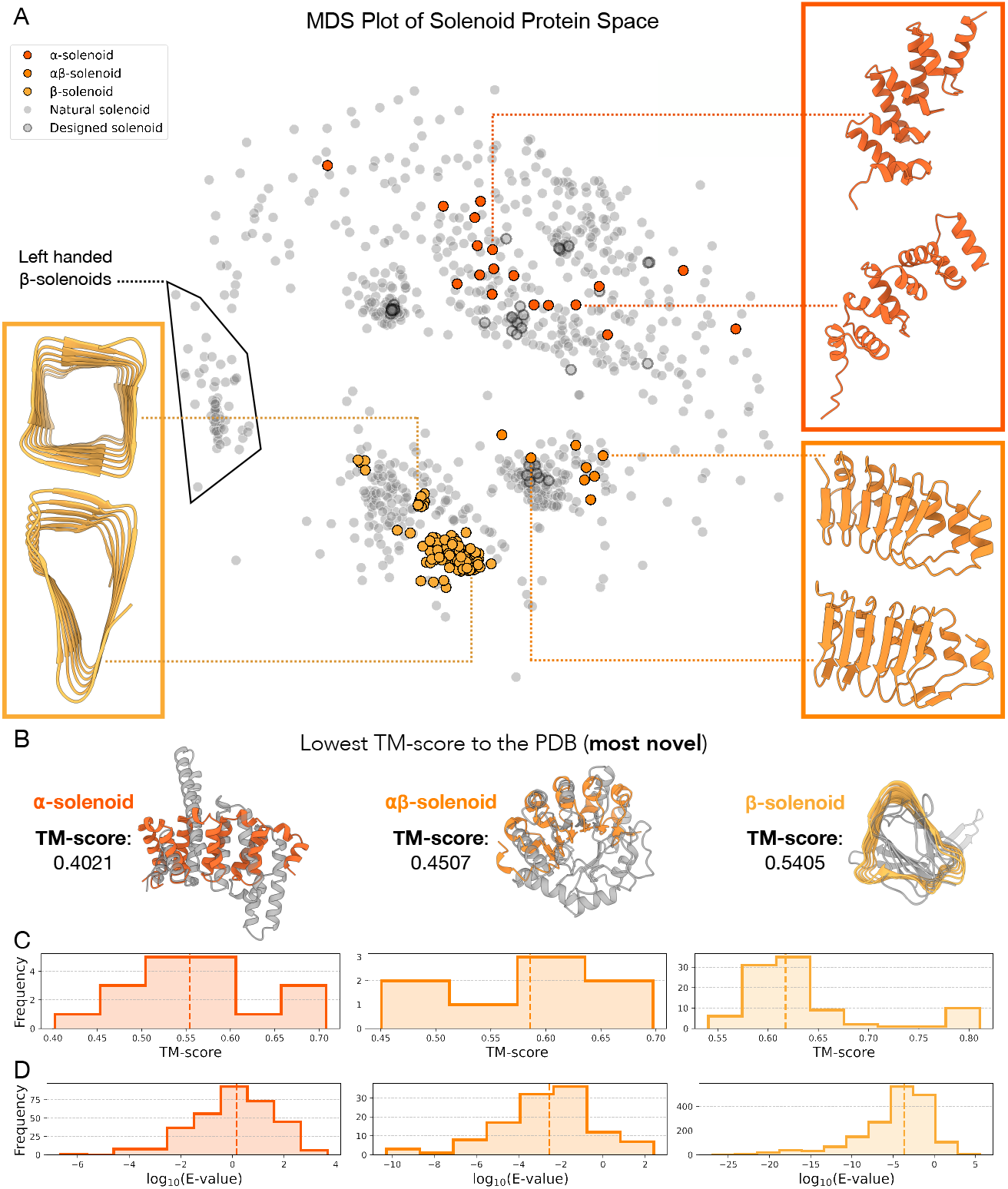
Designed solenoid protein novelty and diversity. **(A)** Natural and novel solenoid protein representation space structural similarity quantified by MDS of solenoid pairwise structural similarities. The solenoids produced by the in silico evolution loop are seen in color. Natural solenoid proteins are in grey and previous designed proteins in the PDB are grey with a black outline. Left-handed β-solenoids are outlined. **(B)** For each solenoid design, we identified the most similar structure in the PDB. Shown here are the least similar of these best matches, the alignments with the lowest TM-scores, for each solenoid type. These represent the most structurally novel designs. The lowest best-match TM-scores were: α-solenoids, 0.40 (PDB:2O8P); αβ-solenoids, 0.45 (PDB:4I6K); β-solenoids, 0.54 (PDB:1K8F). **(C)** Histograms showing the TM-score of the most similar structure in the PDB to the solenoid backbone designs for α-solenoids, αβ-solenoids and β-solenoids. Median TM-scores (dotted line) for the α-, αβ-and β-solenoids are 0.55, 0.59 and 0.62, respectively. **(D)** Histograms showing the E-value distribution for the closest hit in the non-redundant database for ProteinMPNN re-designed sequences, based on backbones generated from the in silico evolution loop for α-solenoids, αβ-solenoids, and β-solenoids. Only sequences with an RMSD <5 Å compared to the original backbone after AF2 prediction, and a pLDDT >80, are included. The median E-values (dotted line) for the α-, αβ-, and β-solenoids are 1.44, 2.92 × 10^−3^, and 2.13 × 10^−4^, respectively.

Among the solenoid types, α-solenoids had the highest structural diversity (**Fig. 2a**). This is likely due to the flexibility in how helices can be paired and arranged, allowing for a broader range of configurations. This observation is consistent with trends seen in natural solenoid proteins, where α-solenoids have the most diverse range of subclassifications and variability [29].

Although natural β-solenoids are either left- or right-handed relative to the solenoid helical axis, with an approximate 70/30 split in the PDB [30], all designed β-solenoids generated by in silico evolution thus far have been exclusively right-handed (**Fig. 2a**). This bias toward right-handedness could stem from the early generations of the evolution trajectories. During these early stages, low-quality output structures with poor pLDDT scores adopt a right-handed β-solenoid, even in cases where different solenoid types were selected for (**Fig. 1c, Supplementary Fig. 1**). Once this handedness is established, it becomes challenging to reverse, as changing handedness is a significant structural change, ultimately steering the process toward high-quality right-handed β-solenoids. A similar trend is observed in the AlphaFold Database, where many low-confidence structures also adopt right-handed β-solenoid-like shapes [31], which suggests AF2 could be biased towards right-handed β-solenoids for certain repetitive sequences. In contrast, the α- and αβ-solenoids have a mixture of left- and right-handed helices, mirroring the diversity observed in natural solenoids.

To quantitatively assess structural novelty, designed solenoids were compared to their closest structural matches in the PDB using Foldseek [32] (**Fig. 2b, c**). TM-score was used as the similarity metric, ranging from 0 (completely different) to 1 (identical). In classification systems such as SCOP and CATH, TMscores below 0.5 typically indicate distinct folds [33]. Based on this threshold, 22% of α-solenoids and 25% of αβ-solenoids were classified as structurally novel (**Fig. 2c**). In contrast, no β-solenoids met this threshold, indicating they all closely resembled existing folds (**Fig. 2c**).

The sequence novelty of ProteinMPNN re-designed solenoids was assessed using MMseqs2 [34] to identify the closest hits in the non-redundant database (nr) [35] (**Fig. 2d**). The E-value distributions were used to evaluate similarity, with lower E-values indicating greater novelty. A threshold of E-value less than 10^−5^ was considered novel [36]. Only sequences consistent with the original design after AF2 reprediction were included, meeting criteria of RMSD less than 5 Å compared to the original backbone and pLDDT greater than 80. Using the E-value cutoff for novelty, 99.7% of α-solenoid, 84.5% of αβ-solenoid, and 64.3% of β-solenoid sequences were classified as novel (**Fig. 2d**). Although no novel backbone structures were identified for the designed β-solenoids (**Fig. 2c**), their backbones still support sequences that have no similarity to natural sequences. This trend is seen across all the designed solenoids, where sequences tend to be more novel than their corresponding structures, as expected given the 3 to 10 times greater conservation of structure over sequence [37].

### Biophysical characterisation of designed solenoids

We selected 24 solenoid designs for experimental characterisation, filtering for backbones with less than 5 Å RMSD to the original design and pLDDT more than 80 across all five AF2 output models (**Supplementary Fig. 2, Supplementary Table 1**). These included 10 backbone designs (3 α-, 3 αβ-, and 4 β-solenoids), with 3-4 redesigned sequences per backbone.

Of the 24 designs, 22 (92%) were successfully cloned into an expression vector, and 9/22 (41%) produced soluble protein at the expected molecular weight following small-scale expression and Ni-NTA purification. Expression and purification success rates were 100% and 100% for α-solenoids, 67% and 50% for αβ-solenoids, and 45% and 9% for β-solenoids respectively. The 9 soluble designs (5 α-, 3 αβ-, and 1 β-solenoid) were taken forward to large scale expression and purification with Ni-NTA and size exclusion chromatography (SEC). All α-solenoids had a major monomer peak at the expected elution volume (**Fig. 3a**), while αβ- and β-solenoids eluted mainly in the void volume or with no clear peak at the expected elution volume (**Supplementary Fig. 3**). Circular Dichroism (CD) spectroscopy was consistent with α-helical secondary structure for all α-solenoids, and thermal stability assays showed that all five α-solenoids retained their secondary structure up to 90°C (**Fig. 3a**).

**Figure 3.**
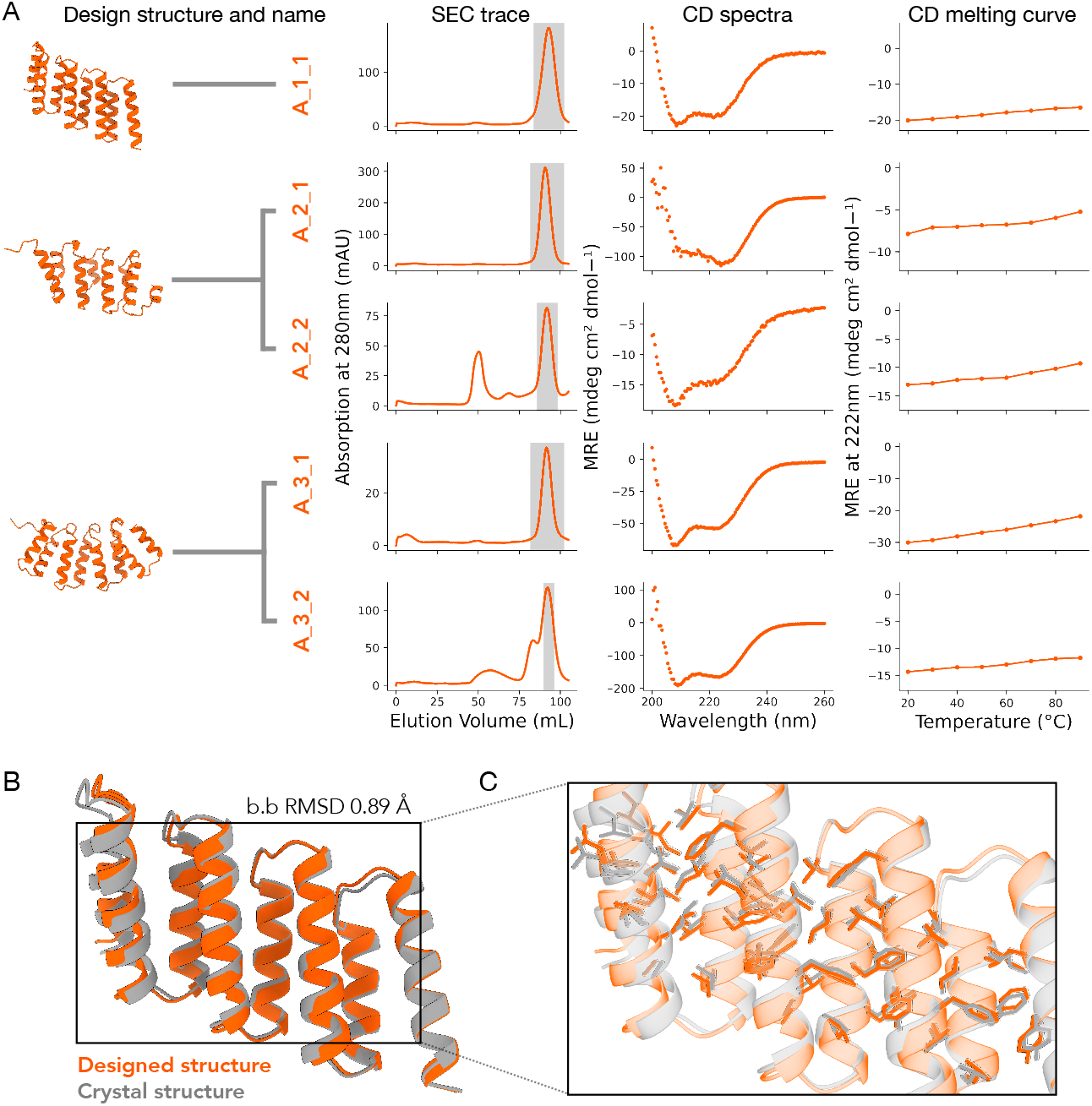
Experimental characterisation of designed solenoid proteins. **(A)** First panel, predicted structure and name of the design. Multiple designs with different sequences but the same protein backbone are grouped under the same backbone structure. Second panel, SEC chromatogram. The highlighted region indicates the fractions pooled to continue for further structural characterisation. Third panel, CD spectra taken at 20°C. Fourth panel, 222 nm CD thermal scan from 20°C to 90°C. **(B)** Superposed designed model (*orange*) and crystal structure (*grey*) of design A_1_1. **(C)** Detail overlay of the crystal structure (*grey*) and designed model (*orange*) with selected hydrophobic core side chains shown as sticks.

To evaluate design accuracy, we attempted crystallisation of the 5 successful α-solenoid designs (**Fig. 3a**). We solved the X-ray crystal structure for A_1_1 at a resolution of 2.8 Å (**Fig. 3b, c, Supplementary Table 2**). The structure showed strong agreement with the design model, with a Cα backbone RMSD of 0.89 Å. The closest structural match in the PDB to A_1_1 was an RNA-binding HEAT repeat protein in humans (PDB: 8BFH), with a TM-score of 0.55 [38]. The A_1_1 sequence showed no significant similarity to known sequences in the nr database, with a lowest E-value of 73.54. To our knowledge, this is the first experimentally determined structure of an octotricopeptide (38mer repeat) α-solenoid. While such repeat architectures occur in nature, no structures of this kind have previously been deposited in the PDB [39].

In summary, all cloned α-solenoid designs expressed, adopted the correct secondary structure, and were thermostable. In contrast, αβ- and β-solenoids failed to produce stable, well-folded proteins. The success rates were 5/6 for α-solenoids, 0/6 for αβ-solenoids, and 0/12 for β-solenoids, with experimental success defined here as a clear SEC peak at the expected elution volume and a CD trace consistent with the designed secondary structure. Despite doubling the number of β-solenoid designs tested, none were viable, highlighting the difficulty in designing β-majority proteins, which often face challenges such as aggregation, misfolding, and insolubility [26, 40].

### Evaluating metrics for predicting experimental outcomes

Design filtering is a major bottleneck in de novo protein design, as successful candidates are often hidden among many designs that will fail at some stage of experimental validation. The goal of filtering is to use computational metrics that can score and select designs with the highest likelihood of experimental success. The use of AF2 in design filtering has significantly improved experimental outcomes by enabling more accurate assessments of structure [14, 41]. However, despite achieving high AF2 confidence scores and low RMSD to the original designed backbones, the first round of αβ- and β-solenoids designs performed poorly experimentally (**Fig. 4a**). This highlighted the need for refining filtering criteria and incorporating alternative metrics. The next round of solenoid designs only included β-solenoids, making the refinement of filtering strategies for β-strand majority proteins a priority.

**Figure 4.**
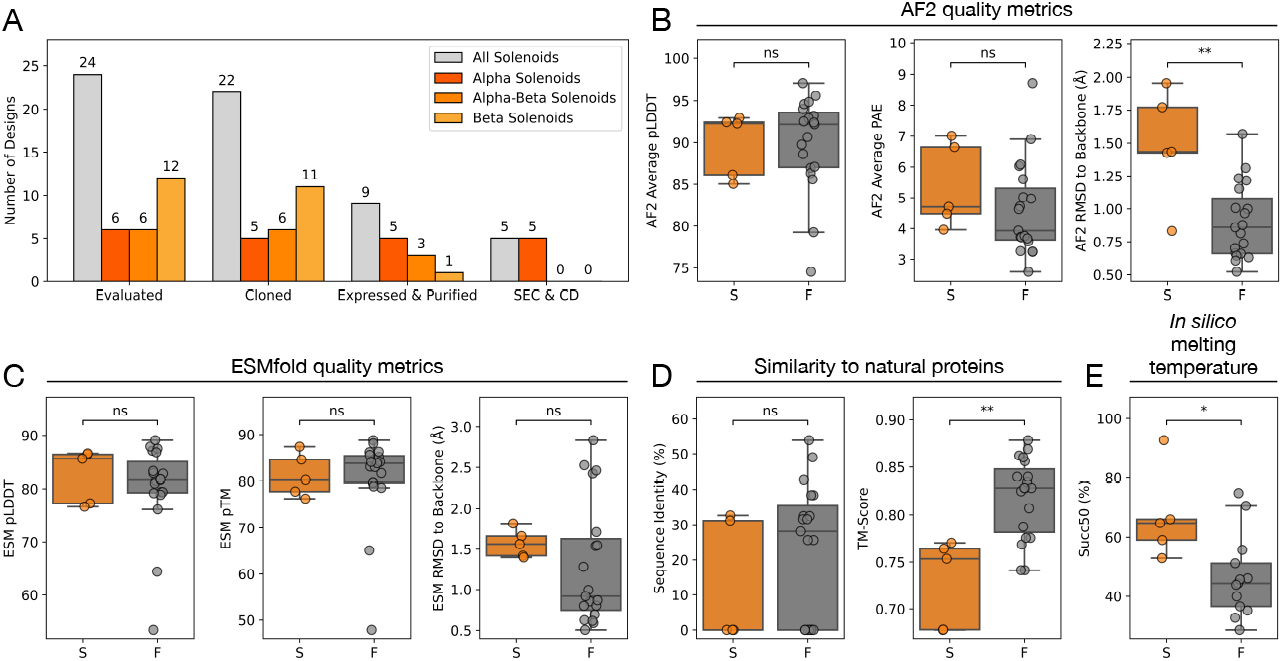
Comparing in silico metrics with experimental outcomes for solenoid designs from the in silico evolution platform. **(A)** bar chart showing the outcomes of the experimental characterisation of *in silico* platform solenoid designs. This is shown by how many designs were evaluated, cloned into a backbone vector, had a purification product at the expected molecular weight after Ni-NTA purification and if there is an expected SEC and CD spectra trace. **(B-E)** Metrics are comparing experimental success (S) versus failure (F) in designs, defined by the presence of an expected SEC peak and a matching CD trace for secondary structure. The five successful designs include A_1_1, A_2_1, A_2_2, A_3_1 and A_3_2; the remaining 19 designs are classified as failures. **(B)** AF2 quality metrics: Box plots for average pLDDT (left, not significant), average PAE (middle, not significant), and RMSD to the original backbone design post-sequence redesign (right, significant with P-value 0.0093). **(C)** ESMFold quality metric*s*: Box plots for pLDDT (left, not significant), pTM (middle, not significant), and RMSD to the original backbone design post-sequence redesign (right, not significant). **(D)** Similarity to natural proteins: Sequence identity with the top hit in the nr database (left, not significant) and TM alignment score to the closest PDB match (right, significant with P-value 0.0044). **(E)** Box plots of the sigmoidal inflection point values (representing ‘melting temperatures’) of the ‘melting curves’ (significant with P-value 0.019). Statistical significance was determined using the Mann-Whitney U test.

To identify better filtering metrics, we conducted a retrospective analysis of solenoid designs, derived from the in silico evolution pipeline, that had previously been experimentally tested. We compared experimentally successful and failed designs using three categories of metrics: AF2 quality metrics (**Fig. 4b**), ESMFold quality metrics (**Fig. 4c**), and similarity of designs to natural proteins (**Fig. 4d**).

Among the original filtering criteria, only two metrics were statistically significant: the RMSD between the original backbone and AF2-repredicted structures, where higher RMSD correlated with successful designs (P-value: 0.0093), and the TM-score to the closest structure in the PDB, where greater structural dissimilarity to natural proteins was linked to experimental success (P-value: 0.0044). These results were counterintuitive, lower RMSD values to the original backbone are usually considered an indicator of design accuracy [14, 17], and recapitulating natural protein structures has a higher probability of success. However, the median RMSD difference of 0.57 Å between the original backbone and AF2-repredicted structures, and the TM-score difference of 0.07 to the closest match in the PDB between successful and failed designs, are both small (**Fig. 4b, d**).

While structural quality metrics such as AF2 confidence scores and RMSD thresholds were sufficient to filter out very poorly performing designs, they failed to distinguish false positives among the remaining high-confidence candidates. Given these results, we explored whether alternative in silico metrics could better predict the experimental outcomes of these designs. ESMFold-based in silico melting has previously been shown to effectively distinguish folded and misfolded designs in β-barrel proteins [42], another β-strand majority protein type. We hypothesised that this approach could also be applied to β-solenoid designs. In silico melting evaluates robustness to sequence perturbation by calculating “melting temperatures” (Succ50) by increasing sequence masking applied to the language model of ESMFold and observing the quality of output predictions [42].

Applying this method to our experimentally characterised solenoids, we observed a statistically significant correlation between higher Succ50 values and experimental success (**Fig. 4e, Supplementary Fig. 4**). The median Succ50 difference was 20.4% masking (P-value: 0.019), with higher predicted melting temperatures linked to successful designs. This aligns with expectations, as higher Succ50 values indicate greater robustness to in silico sequence perturbation and, therefore, a potentially more robust structure.

The synthetic solenoid SynRFR24.1, which was designed outside of our pipeline [43], had a high Succ50 value of 71%, significantly higher than all our β-solenoid designs (**Fig. 4e**). This shows that the observed separation is not simply due to β-solenoids inherently having lower Succ50 values compared to α-solenoids. Furthermore, the crystallised α-solenoid from our pipeline had the highest Succ50 value of 93%, far outperforming all other designs in the round (**Fig. 4e**).

### Design and experimental validation of capped β-solenoids

None of the initial β-solenoid designs from the in silico evolution platform were experimentally successful. Unlike α-solenoids, which naturally terminate with stable repeating units that act as effective caps, β-solenoids in Nature often require dedicated terminal capping regions [44, 45]. While ProteinMPNN can introduce terminal regions by incorporating amino acids that act as capping elements for the final repeat, the large hydrophobic cross-section of β-solenoids makes sequence-level redesign insufficient to fully address the issue. Therefore, the β-solenoids from the initial experimental round could have lacked proper capping and had exposed hydrophobic cores. We hypothesised that designing terminal capping regions for β-solenoids, rather than relying solely on sequence redesign, could improve the experimental performance of these designs.

The β-solenoid backbones generated through the in silico evolution loop were used as scaffolds to design N- and C-terminal capping regions using ProteinGenerator [46] (**Fig. 5a**). Capping regions were introduced across a set of 94 β-solenoid backbones. Capped designs were classified as “designable” if they met the following criteria: less than 2 Å RMSD to the backbone design, less than 5 Å pAE for the whole structure, less than 5 Å pAE for the terminal regions, and more than 80 pLDDT from both AF2 and ESMFold. Using these thresholds, we generated 2,374 capped designs for 5mer repeats, 10,787 for 25mer repeats, and 3,975 for 30mer repeats.

**Figure 5.**
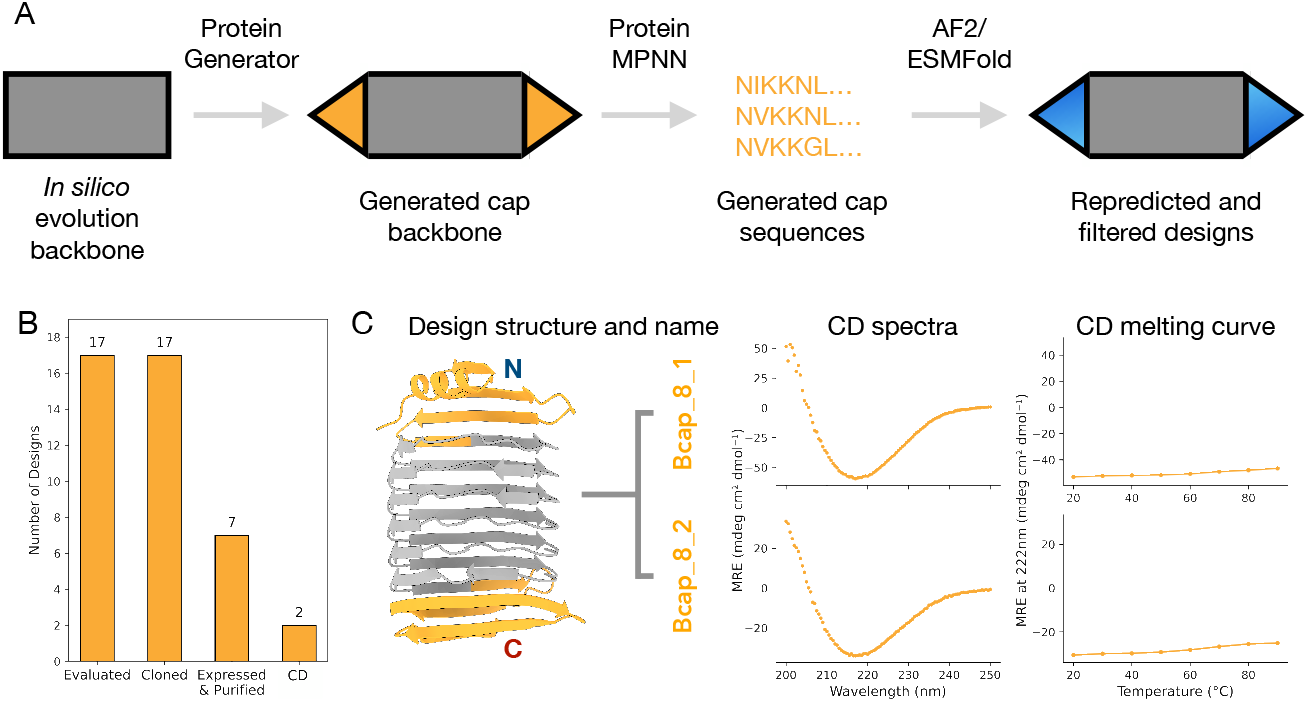
Design and experimental characterisation of capped β-solenoids. **(A)** Pipeline to design capping regions for β-solenoids. **(B)** Outcomes of the experimental characterisation of capped β-solenoid designs. This is shown by how many designs were evaluated, cloned into a backbone vector, had a purification product at the expected molecular weight after Ni-NTA purification and if there is an expected CD spectra trace. **(C)** First panel, predicted structure and name of the design characterised. β-solenoid region in grey and capping region in yellow. Second panel, CD spectra taken at 20°C. Third panel, CD (222 nm) thermal scan from 20°C to 90°C.

Our first round of β-solenoid designs had shown that structural metrics alone did not correlate well with experimental success (**Fig. 4b, c**). To further refine selection, we applied an in silico melting filter that was shown to correlate with success (**Fig. 4e**). Designs were taken forward only if they met all previous criteria and had a Succ50 value above 71%, a threshold based on the previously designed β-solenoid SynRFR24.1 [43]. To ensure diversity within the selected designs, we used Foldseek [32] clustering based on a TM-score threshold of 0.7. This resulted in 12 distinct clusters, with each representing a different backbone (**Supplementary Fig. 5**).

We selected 17 capped β-solenoid designs for experimental characterisation, filtering for less than 2 Å RMSD to the backbone design, less than 5 Å pAE for the overall structure and terminal regions, more than 80 pLDDT from both AF2 and ESMFold, and a Succ50 value above 71% (**Supplementary Fig. 5, Supplementary Table 3**). These included a selection of 8 different backbones, with 4-1 redesigned sequences per backbone.

All constructs (100%) were successfully cloned into a backbone vector. Large-scale expression, followed by Ni-NTA purification, showed that 7/17 designs (41%) produced soluble protein at the expected molecular weight (**Fig. 5b**). The 7 soluble capped β-solenoid designs were taken forward for further purification and biophysical characterisation. Of these, Bcap_8_1 and Bcap_8_2 showed elution peaks in anion exchange chromatography (AEC) following SEC that corresponded to the expected band on SDS-PAGE (**Supplementary Fig. 6**). CD spectroscopy of the corresponding AEC fractions confirmed that the designs Bcap_8_1 and Bcap_8_2 had β-strand secondary structure (**Fig. 5c**). We assayed the thermal stability of Bcap_8_1 and Bcap_8_2 and both maintained their secondary structure up to 90°C (**Fig. 5c**). While we were able to crystallise Bcap_8_2, the crystals only diffracted to 10 Å resolution and the structure could not be solved. However the ability of the protein to crystallise provides supporting evidence that it is folded (**Supplementary Fig. 7, Supplementary Table 4**).

The introduction of capping regions and addition of new filtering metrics improved the success rate of β-solenoid designs with biophysical characteristics supporting their designed structure, increasing from 0% (0/12) in the initial uncapped designs to 12% (2/17) for capped β-solenoid designs (**Fig. 5b**). Beyond terminal aggregation, β-solenoids face intrinsic folding challenges. β-strands are susceptible to misfolding due to their extensive non-local interactions, which slow down folding and increase the likelihood of aggregation [26]. The edges of β-sheets are sticky sites that can engage in aggregation and amyloid formation. These factors could have contributed to the poor performance of β-solenoids relative to α-solenoids.

Despite high AF2 confidence scores, the β-solenoid designs may have failed experimentally because the backbone structures themselves were undesignable. AF2 can hallucinate high-confidence β-solenoid structures for repetitive sequences, generating implausible features that do not correspond to physically stable proteins [31]. Additionally, β-solenoid designs converged quickly during the in silico evolution pipeline, often within as few as five generations, this was significantly faster than α- or αβ-solenoids. While this initially appeared promising, it may indicate that the design search space is prematurely collapsing onto experimentally non-viable solutions.

## Discussion

A central goal of de novo protein design is to create proteins with specific structural and functional properties. Recent advances in deep learning, particularly AF2 [11], have revolutionised protein design by providing unprecedented insights into sequence-structure relationships. Here, we present an in silico evolution platform that integrates AF2 as an oracle with a genetic algorithm to hallucinate novel α-, β-, and αβ-solenoid proteins. This approach enables the exploration of both natural and novel solenoid structures and is the first hallucination method that we are aware of to generate specific folds from scratch, independent of natural sequence or structural starting points, by using a discriminator network. While the in silico evolution platform uses a solenoid fold-specific discriminator, the module could be replaced with a classifier for any chosen fold, allowing for the generation of diverse proteins with specific folds.

A critical question in generative protein design is how novel and diverse the resulting structures are. The solenoid designs spanned both known and novel solenoid structures, with α-solenoids having the highest structural diversity. In contrast, all β-solenoid designs were exclusively right-handed, despite the PDB containing both right- and left-handed β-solenoids. This raises the question of whether AF2 has a bias towards right-handed β-solenoid structures for highly repetitive sequences, a trend also observed in the AlphaFold Database which has many instances of right-handed implausible β-solenoid structures [31]. While AF2 is known to produce artifacts such as highly confident isolated long helices, the bias towards right-handed β-solenoids is unusual, as structured domains with extensive inter-residue contacts are generally reliably predicted [2]. By repurposing AF2 for design, we highlight its tendency to favour right-handed β-solenoid structures for certain highly repetitive sequences.

Using the in silico evolution platform, we generated a library of 121 solenoid protein backbones, selecting 10 designs for experimental characterisation. Out of the 10 backbone designs, 6/10 showed soluble expression and purification products and 3/10 showed SEC elution peaks and CD spectra consistent with their designed structures, with one α-solenoid design further validated by a crystal structure closely matching the model.

However, all of the experimentally successful solenoid proteins were α-solenoids, while all β-solenoids initially failed despite testing twice the number of constructs experimentally, reflecting the difficulty of designing β-strand majority proteins. We reasoned this disparity could be due to three key factors: aggregation from the exposed edges of β-sheets and exposed hydrophobics in the terminal regions of β-solenoids, the inherently slow folding kinetics of β-majority folds due to their reliance on non-local interactions, which increases the likelihood of misfolding and aggregation, and adversarial backbone and sequence design when using AF2 as both an oracle and a post-design filter.

By introducing terminal capping regions and refining designs based on earlier experimental screens, we stabilized two β-solenoid constructs that had CD spectra consistent with folded β-solenoids. These results highlight the value of pairing computational generation with iterative laboratory feedback to address problems such as misfolding and aggregation. Despite this improvement, β-solenoids still performed significantly worse than α-solenoids, which consistently had expected biophysical characteristics and good yields. This suggests that aggregation is not solely due to exposed terminal regions but is also influenced by factors such as folding stability. AF2 can also give incorrectly confident predictions of β-solenoid structures for sequence repeats, which might be considered adversarial inputs [31].

A major distinction between hallucination and diffusion-based generative methods is how they explore design space. Diffusion samples from a learned distribution and requires extensive filtering post hoc, discarding a high proportion of candidates. Hallucination, as in the in silico evolution platform presented here, optimizes sequences directly on AF2, which resulted in a 99% designability rate for our solenoid designs. However, even with direct optimization, filtering remains critical for identifying experimentally viable proteins. Our results suggest that standard structure predictor confidence metrics effectively eliminate poor candidates but fail to distinguish false positives. By refining our filtering strategy based on experimental outcomes and incorporating in silico melting, we improved success rates. This highlights the need for additional filtering metrics to complement AF2 confidence scores, providing a more reliable way to identify viable designs. Until in silico design methods become much improved, we will still require these filtering metrics.

Beyond developing an experimentally validated hallucination-based approach for generating fold-specific proteins, this work addressed two key challenges in protein design: optimising filtering metrics to improve design selection and designing β-stand majority proteins. Our results demonstrate that hallucination-based design can generate fold-specific solenoid proteins de novo with high designability and experimental success, particularly for α-solenoids. With improved filtering and iterative feedback, this approach offers a promising route to expanding the repertoire of stable, functional de novo proteins.

## Methods

### in silico evolution platform

We developed an in silico evolution platform using AF2 as an oracle for protein design (**Fig. 1a**). This is similar to work by Anishchenko *et al*. [15], who used trRosetta for whole-protein hallucination and more recent hallucination pipelines that use AF2 and MCMC for optimisation [17, 21, 47, 48]. The pipeline has two main components: a sequence optimiser and a structure predictor.

The sequence optimiser is a genetic algorithm based on a greedy search method for biological sequence design [49]. We use a Wright-Fisher selection scheme, where higher-scoring parents have an exponentially higher probability of contributing offspring. In each generation, we create new children by randomly mutating each position in a parent sequence with probability 1/L, where L is the sequence repeat length. On average, this yields one mutation per offspring, we reasoned this was suitable due to the small repeat lengths being sampled. We use the 20 canonical amino acids and choose a new amino acid uniformly from the remaining 19 at any mutated site. Child sequences that are distinct and score well replace the worst-performing members of the current population in a “steady-state” fashion, thereby favoring incremental improvements in each round. Once a sequence meets the design threshold criteria, it is returned as a complete design, and the optimisation loop stops. Runs failing to reach the threshold within 30 generations were terminated.

Structure prediction was performed using Colabfold [50] locally in single-sequence mode, with the number of recycles set to 1. All five prediction models were averaged for pLDDT and solenoid scores. To pass the design threshold, all five models were required to exceed a minimum pLDDT of 70.

We normalized the pLDDT score to 0-1, derived a 0-1 solenoid classification probability from SOLeNNoID [24], and combined them with equal weight to yield a single cost measure on the same 0-1 scale. The detector classified proteins as β-solenoid, α-solenoid, αβ-solenoid, or non-solenoid and assigned per residue probabilities for each classification. These probabilities were converted to one-hot encoded classifications, summed, and averaged across the structure to evaluate solenoid likelihood.

The type of solenoid protein produced in each trajectory was guided by the initial conditions: repeat length (L), number of repeats (N), solenoid category score from the solenoid discriminator, and thresholds for the solenoid score and pLDDT in the cost function. Repeat lengths were chosen based on ranges typical of natural solenoid proteins [51-53]. The number of repeats was adjusted to ensure scaffolds contained at least four repeating units, resulting in proteins ranging from 130 to 180 amino acids in length.

The sequences from the in silico evolution loop were re-designed using ProteinMPNN [54]. For each protein backbone, 20 sequences were generated with ProteinMPNN at a temperature of 0.2, disallowing cysteines. The redesigned sequences were repredicted with AF2, and the resulting structures were ranked based on the pLDDT and RMSD relative to the original backbone. Only structures with less than 5 Å RMSD to the original backbone and more than 80 average pLDDT across all five models were considered suitable for experimental characterisation.

### β-solenoid capping design pipeline

We developed a pipeline for adding capping regions to β-solenoid proteins (**Fig. 5a**). This involved three stages: backbone designability testing, N-terminal cap design, and C-terminal cap design.

In the backbone designability stage, β-solenoid backbones generated through the in silico evolution loop were evaluated for their “designability”. For each backbone, 10 redesigned sequences were generated using ProteinMPNN at a temperature of 0.2, disallowing cysteines [54] and re-predicted with ColabFold [50] in single-sequence mode and ESMFold [55]. Backbones were considered “designable” if they passed filtering thresholds of less than Å RMSD to the backbone, less than 5 Å pAE from AF2 overall, and more than 80 pLDDT from AF2 and ESMFold. These criteria were applied across all filtering stages of the pipeline.

ProteinGenerator [46] was then used to create the capping regions for the β-solenoids. For the N-terminal cap design, ProteinGenerator was used to generate 10 N-terminal cap structures, and ProteinMPNN produced 10 corresponding sequences for these structures. These sequences were re-predicted and filtered using the same structural criteria established in the backbone designability stage including an additional less than 5 Å pAE from AF2 for the capping region alone. For successful candidates with high-quality N-terminal caps, the process was repeated to design C-terminal caps, using the capped N-terminal structures as starting points.

ProteinGenerator was configured to exclude cysteine residues and specify a repeating secondary structure pattern of “HXX” (H = α-helix and X = any structure) to introduce a helical bias for the caps without enforcing exclusively helical regions, which otherwise resulted in single long α-helices. The length of the capping region used was twice the repeat length for most designs. For shorter 5mer repeats, the cap length was extended to four times the repeat length.

### In silico melting experiments

The in silico melting experiments were performed based on the method described by Hermosilla *et al*. (2024) [42]. ESMFold [55] was used to generate “melting curves” for each protein design by masking random positions in the input sequence to the language model. Masking levels ranged from 30% to 99%, applied in 5% increments. At each masking level, 64 models were generated with randomly applied masking patterns. The success rate at each masking level was defined as the fraction of the 64 models with a predicted TM-score (pTM) greater than 0.75.

The success rate as a function of the masking level was then fitted to a sigmoidal curve to model the “melting” behaviour of each design. The inflection point of the fitted sigmoidal curve (Succ50), referred to as the melting point, was extracted as a measure of robustness to sequence perturbation. Designs that could not be fitted to a sigmoid curve were excluded from the analysis. Additionally, melting points outside the 0-1 range were discarded as unfeasible.

### PDB solenoid database

Annotated natural β-, α- and αβ-solenoid protein families were obtained from the RepeatsDB database [56]. The database was prepared using a similar approach used by Brunette *et al*. (2015) for their repeat protein analysis [8]. A Python script was developed to download PDB coordinates, extract relevant protein chains, and retrieve details such as repeat region boundaries, average repeat unit length, starting residue indices for repeats, and insertion sites. Redundancy was removed using CD-HIT, applying an all-against-all sequence clustering at 90% identity [57]. A single representative sequence was selected for each cluster, and only the solenoid region was used. In total, 495 α-, 164 αβ-, and 209 β-solenoids from PDB structures were used for comparison against hallucinated solenoid designs.

### Building solenoid structure representation space

A visualisation of the structural space with both natural and novel solenoids was created using the curated PDB solenoid database and the hallucinated solenoid structures. Previously designed solenoid proteins were identified using SOLeNNoID [24] to detect solenoids within data from the Protein Design Archive [58]. An all-against-all structural alignment for the hallucinated solenoids against the solenoids in the PDB was run with Foldseek [32]. A Python script converted the pairwise TM-scores from Foldseek into a pairwise distance matrix, which was then embedded into 2D space using MDS with the scikit-learn library [59].

### Structure and sequence comparison

To assess structural similarity, a structural alignment of hallucinated solenoid structures against the PDB was performed using Foldseek [32]. The structures were globally aligned using TMalign mode, with the similarity metric being a TM-score normalised by the query length. For each query, the highest TM-score PDB match was returned. Where clustering was applied, Foldseek was used to cluster hallucinated solenoids with a TM-score threshold of 0.7 [32].

For sequence similarity, designed sequences were searched against the nr protein sequence database using MMseqs [34]. A Python script retrieved the hit with the lowest E-value for each query, representing the most similar natural sequence.

### Codon optimisation for repeat proteins

Despite sequence redesign introducing deviations from perfect repeats, the protein sequences remained highly repetitive. As a result, their corresponding DNA sequences were too repetitive for synthesis using standard *E. coli* codon optimisation alone. A codon optimisation pipeline was developed for highly repetitive DNA sequences based on the DNAchisel library [60].

### Cloning and transformation

Linear DNA fragments encoding the designed protein coding sequences (CDSs) were ordered as a 96-well formatted eBlock from Integrated DNA Technologies. Each sequence was designed with inwards facing flanking SapI restriction sites for Golden-Gate cloning into the pEDEV-HT expression vector. The associated fusion sites were AGT and TAA, allowing in-frame cloning incorporating the serine of the linker and stop codon respectively.

The pEDEV-HT expression vector used for these CDSs (and all subsequent constructs), was a modified pET29b(+) expression vector (Twist Bioscience), containing a bespoke insertion region for Golden Gate cloning and N-terminal tagging. An N-terminal His-tag and thrombin cleavage site were added directly downstream of the original pET29b(+) start codon, followed by an inverted bacterial expression cassette encoding the tsPurple chromoprotein. This cassette was flanked by outward facing SapI sites with AGT and TAA fusion sites such that cloning into this region would result in the excision of the chromoprotein cassette and scarless insertion of the target CDS. This system enabled scarless one-pot-one-step cloning of linear CDSs into the vector, with visual confirmation of successful insertions: colonies carrying the unmodified entry vector appeared purple, while those containing the desired construct appeared white.

Golden Gate assembly reactions were performed using a Biorad T100 thermocycler using an alternating 16°C/37°C ligation/digestion over 30 cycles with two minutes per temperature. Following assembly, the golden gate reaction mixtures were transformed into chemically competent *E. coli* KRX cells (Promega) and incubated overnight at 37°C on kanamycin-supplemented lysogeny broth (LB) agar plates (1.5% agar). Single colonies were selected by color and picked for colony PCR, and the amplicons were analysed by agarose gel electrophoresis to confirm the presence of the correct insert. The same pipette tip used for colony picking was reserved in a Falcon tube for overnight culture if the insert size was verified.

Colonies with the correct insert were grown overnight in 5 mL LB supplemented with 50 μg/mL kanamycin. The overnight cultures were used to make glycerol stocks and a plasmid-miniprep. The glycerol stocks were prepared by 200 µL of 45 % [v/v] Glycerol in LB mixed with 200 µL overnight culture, flash-frozen at -80°C. The remaining overnight culture was plasmid-miniprepped (NEB Monarch Kit) and sent for sequence verification.

### Small scale expression and purification

Small-scale expression screens were conducted to identify successful constructs for further characterisation, including large-scale expression, purification, and downstream analyses. Unless otherwise stated, kanamycin was used at a final concentration of 50 μg/mL.

Glycerol stocks were used to inoculate 5 mL of LB-Kanamycin, which was grown overnight at 37°C with shaking. A 1:100 dilution was prepared by inoculating 5 mL of terrific broth (TB)- Kanamycin with 50 μL of the overnight culture [61]. The cultures were grown for 3.5 hours at 37°C with shaking until they reached an OD600 of 0.8-1. A sample was taken for SDS-PAGE analysis of the uninduced culture. Expression was induced by adding 0.1% [w/v] rhamnose and 1 mM isopropyl β-D-1-thiogalactropyranoside (IPTG), and cultures were incubated for an additional 4 hours at 37°C with shaking. A sample was taken for SDS-PAGE analysis of the induced culture.

The cultures were transferred into a 96-well plate, with each sample in a separate well, for a final volume of 2 mL per well. The plate was centrifuged at 4,000 × g for 5 minutes at 4°C, and the growth medium discarded by inverting the plate. The harvested cell pellets were then frozen at -20°C in the plate.

Frozen pellets were thawed and resuspended in 300 µL of B-PER chemical lysis buffer (Thermo Fisher Scientific) supplemented with 0.1 mg/mL lysozyme (from a 100 mg/mL stock in 50% [v/v] glycerol), 25 units of DNase I (New England Biolabs), 1 mM PMSF (from a 100 mM stock in propan-2-ol), and an EDTA-free protease inhibitor tablet (Roche). The plate was sealed with a microplate seal, vortexed for 5 minutes until fully resuspended, and incubated at room temperature for 15 minutes. The lysate was centrifuged at 4,000 rpm for 20 minutes at 4°C, and a sample was taken for SDS-PAGE analysis of the clear lysate.

Protein purification was performed using His-tag-based immobilized metal affinity chromatography. Ni-NTA resin was prepared by adding 150 µL of Ni-NTA slurry to each well of a 96-well fritted plate (Pall AcroPrep 8130), for a final bed volume of 75 µL after centrifugation at 3,000 × g for 5 minutes at 4°C. The same collection plate was used for all purification steps, with supernatants either collected or discarded at each stage. The Ni-NTA resin was equilibrated with 3 × 200 µL wash buffer (20 mM Tris, 150 mM NaCl, 25 mM imidazole, pH 8.0), each followed by centrifugation at 3,000 × g for 5 minutes at 4°C.

After equilibration, 200 µL of the clear lysate was applied to the resin and centrifuged at 3,000 × g for 5 minutes at 4°C. The resin was washed with 2 × 200 µL wash buffer, each step followed by centrifugation at 3,000 × g for 5 minutes at 4°C, and a sample was taken for SDS-PAGE analysis of the wash fractions. Proteins were eluted by adding 200 µL of elution buffer (20 mM Tris, 150 mM NaCl, 500 mM imidazole, pH 8.0) to each well, incubating for 5 minutes, and eluting by centrifugation at 3,000 × g for 5 minutes. A sample was taken for SDS-PAGE analysis of the eluate, and the remaining protein was stored at 4°C for further processing.

### Large scale expression and purification

Glycerol stocks were used to inoculate a 10 ml LB-Kanamycin overnight starter culture, grown at 37°C with shaking. The starter culture was used to inoculate 1 L of TB medium, which was grown to an OD600 of 0.8-1 at 37 °C before inducing expression with 0.1% (w/v) rhamnose and 1 mM IPTG. Cultures were incubated overnight at 18°C with shaking at 200 rpm.

Each 1 L culture was harvested by centrifugation at 4,000 rpm for 30 minutes, and the resulting cell pellet was resuspended in 30 mL of lysis buffer (20 mM Tris, 150 mM NaCl, pH 8.0) supplemented with an EDTA-free protease inhibitor tablet (Roche). Cells were lysed by passing the suspension through a cell disruptor at 32 kpsi twice. The lysate was centrifuged at 35,000 rpm for 45 minutes at 4°C to pellet cellular debris, and the soluble fraction (clear lysate) was incubated with 1 mL of Ni-NTA beads overnight at 4°C with gentle rotation.

Batch purification was used for the Ni-NTA purification stage to accommodate the large sample number, with all steps performed at 4°C. The Ni-NTA bead/soluble fraction was centrifuged at 1,000 x g for 5 minutes, and the supernatant (flow-through) was collected. This was washed with 3 × 15-ml wash buffer (20 mM Tris, 150 mM NaCl, 25 mM Imidazole, pH 8.0), with supernatants collected at each step. Proteins were eluted by washing the beads with 3 × 15 mL volumes of elution buffer (20 mM Tris, 150 mM NaCl, 500 mM imidazole, pH 8.0). Elution fractions were pooled and concentrated.

The combined fractions were further purified by SEC using either a HiLoad 16/600 Superdex 75 pg (120 mL bed volume) or a Superdex 200 Increase 10/300 GL (24 mL bed volume) column on an ÄKTA Pure 25 (GE Healthcare) fast protein liquid chromatography (FPLC) system at 4°C, equilibrated in lysis buffer. Fractions corresponding to major peaks were analysed by SDS-PAGE to confirm the presence of the correct molecular weight species. Verified fractions were pooled and concentrated to 1 mL.

Capped β-solenoid designs were further purified using a HiTrap Q HP anion exchange chromatography column (GE Healthcare) on an ÄKTA Pure 25 FPLC system at 4°C. The proteins were eluted with a linear gradient of 25-500 mM NaCl (in 20 mM Tris, pH 8.0), at a flow rate of 2 mL/min. Fractions corresponding to peaks were verified by SDS-PAGE gels to confirm the presence of the correct molecular weight species. Verified fractions were pooled, buffer-exchanged into lysis buffer and concentrated to 1 mL.

### Circular dichroism

300 µL of sample at 0.2-0.5 mg/mL was prepared for CD analysis. Samples were buffer exchanged from the lysis buffer (20 mM Tris, 150 mM NaCl, pH 8.0) into CD buffer (20 mM Tris, 150 mM NaF, pH 8.0) to prevent chloride interference, which absorbs in the far-UV region.

CD spectra were recorded using an Applied Photophysics Chirascan with samples in 1 mm pathlength quartz cuvettes (Hellma QS Quartz cell). Full spectra were collected from 190-260 nm with a 0.5 nm step size at 20°C. For thermal melts, the temperature was ramped from 20°C to 90°C at 10°C intervals, and a reading at 222 nm taken at each interval.

### Crystallography

Crystallisation trials were performed on selected designs. Proteins were concentrated to 10-50 mg/mL, depending on yield and the number of plates set up, as specified in the results. A range of crystallisation screens was tested to identify suitable conditions for crystal formation. For all screens, protein samples were mixed with the mother liquor in 1:1 (400 nL drop) and 2:1 (300 nL drop) ratio. Crystallisation plates were prepared using a Mosquito robot (TTP Labtech) and incubated at 16°C.

A_1_1 crystallised in 25% PEG 1500, 100 mM SPG buffer (pH 5.5). Crystals were cryoprotected with mother liquor and flash-cooled in liquid nitrogen. Data were collected on beamline I24 at Diamond Light Source. Phaser [62] was used for molecular replacement, using the design model as search model. The structure was built in Coot [63], refined with phenix.refine [64], and validated with MolProbity [65].

Bcap_8_2 crystallised in 0.1 M sodium citrate (pH 5.6) with 35% tert-butanol. Data were also collected on beamline I24 and processed with *xia2* [66] and DIALS [67]. Phaser [62] was used for molecular replacement and tNCS analysis, and *molrep* [68] was used to calculate the self-rotation function.

## Data Availability

The A_1_1 crystal structure and structure factors are deposited in the PDB under accession 9HNH.

## Code Availability

The in silico evolution platform source code is available at https://github.com/dpretorius/InSilicoEvolution.

## Acknowledgements

We thank Sam Horrell and the Imperial Centre for Structural Biology for help with X-ray crystallography. The authors would like to thank Diamond Light Source for beamtime (BAG proposal mx31800), and the staff of beamline I24 for assistance with crystal testing and data collection. DP was supported by an EPSRC CDT training grant (EP/S022856/1). GN was supported by an EPSRC DTP training grant (EP/R513052/1). We thank Sidney Lisanza, Dmitri Zorine and Harley Pyles for helpful discussions.

## Author contributions

DP, JWM and GN designed the research program. DP performed the study. KW and SF contributed with protein expression and purification. HT constructed the expression plasmid. DP and SO did the in silico melting experiments. DP and JWM wrote the manuscript with contributions from all authors.

