## Supplementary Figures and Tables for "Designing Novel Solenoid Proteins with In Silico Evolution"

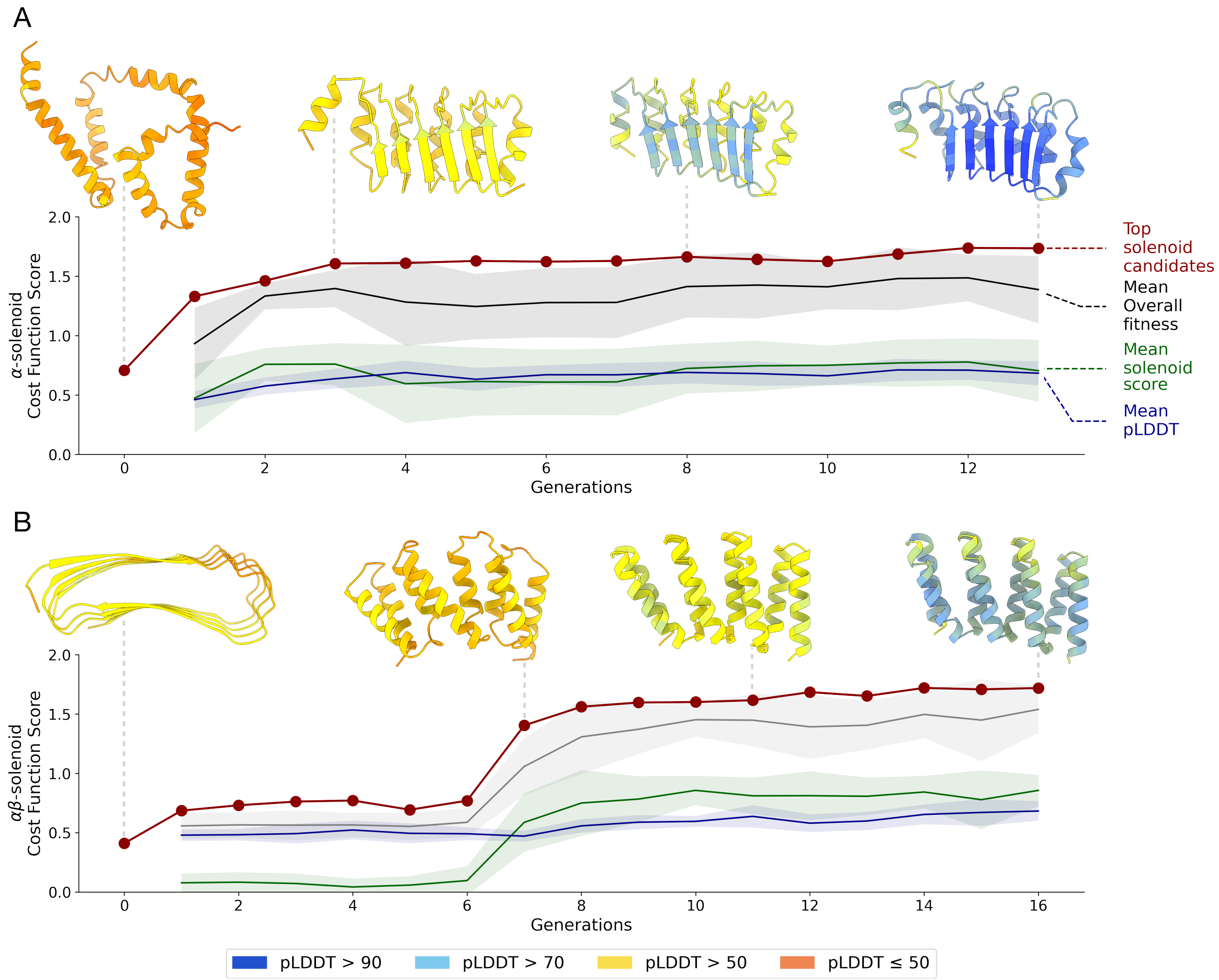


**Supplementary Figure 1 | Solenoid hallucination trajectories.** Examples of progression for **(A)** αβ-solenoid and **(B)** α-solenoid from an initial random sequence to complete solenoid design. Generation 0 is the initial random starting sequence. The cost function value of the best candidate (*red*) is shown for each generation. The average fitness (*black*), pLDDT (*blue*) and solenoid score (*green*) for the whole population of each generation is shown. 1 standard deviation above and below the mean is shown by the shaded region of the same colour. Top candidate structures coloured by pLDDT score are shown for generation **(A)** 0, 3, 8, 13 and **(B)** 0, 7, 11, 16. These were selected to best represent the progression of the structure.

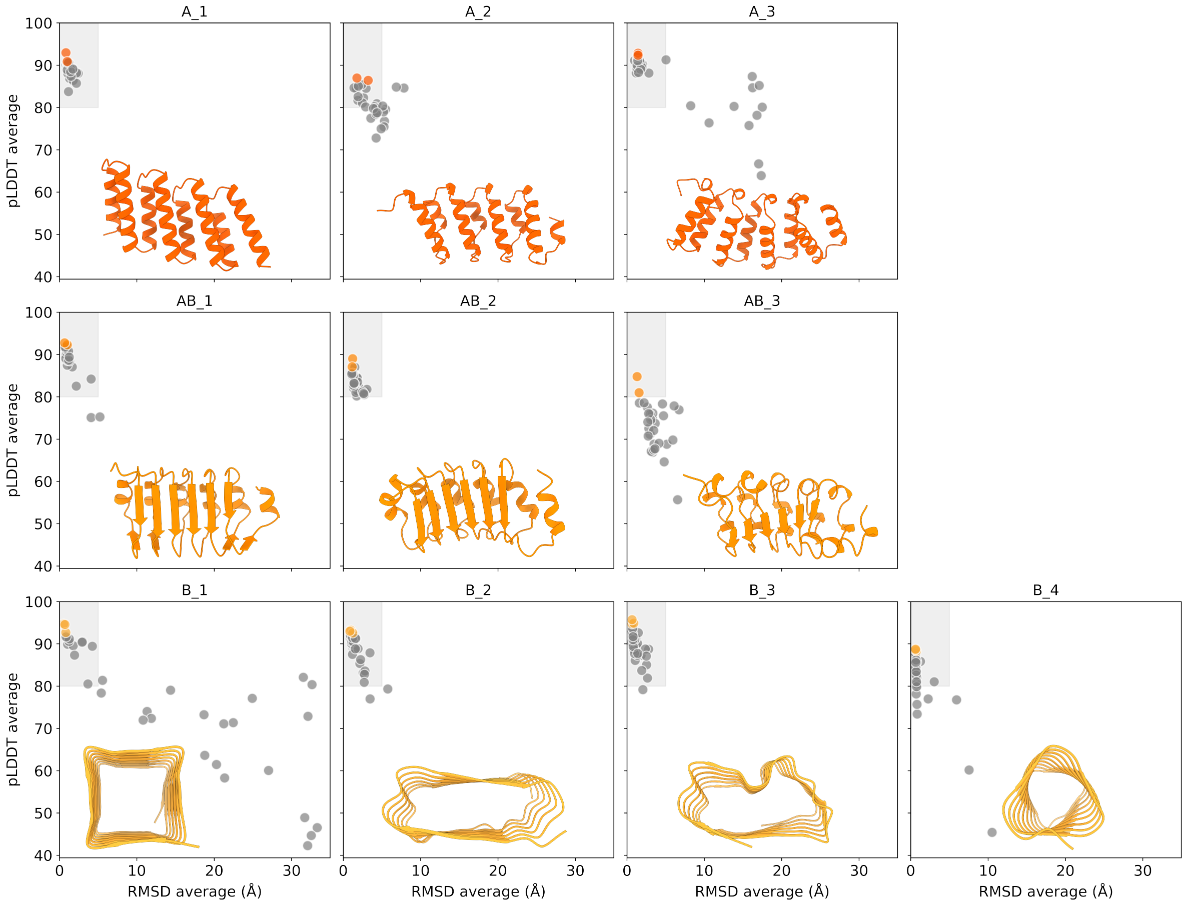


**Supplementary Figure 2 | Quality metrics of solenoid backbones selected for experimental characterisation.** Results from the 20 ProteinMPNN sequence redesign and structure prediction compared to the original backbone for the α-, αβ-, and β-solenoid backbones selected for experimental validation. The top 2 sequences were selected for α-, αβ-solenoids and top 3 sequences for β-solenoids. The grey area indicates the region of design quality which satisfies the requirements of <5 Å RMSD to the original backbone and >80 pLDDT. The backbone structure for each design is shown on the plot.

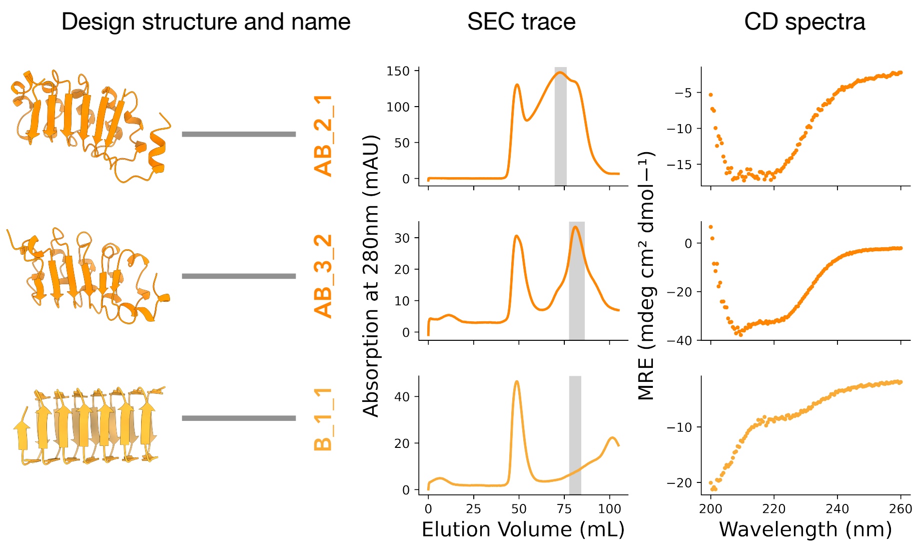


**Supplementary Figure 3 | Biophysical characterisation of unsuccessful large scale solenoid protein expression.** First panel, computational structure and name of the design characterised. Second panel, SEC chromatogram. Highlighted regions indicate the expected monomer elution range, which were pooled for further characterisation. Third panel, CD spectra taken at 20°C.

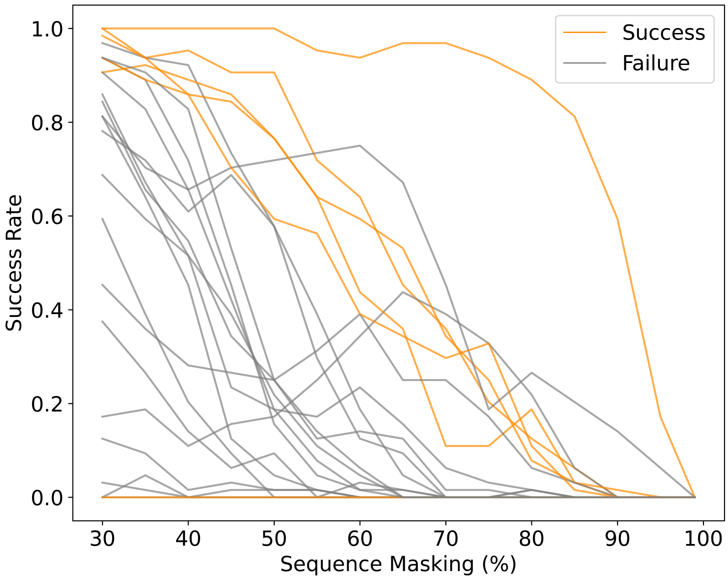


**Supplementary Figure 4 | Comparing in silico masking experiments with experimental outcomes of in silico evolution platform solenoid designs.** Metrics are comparing experimental success versus failure in designs, defined by the presence of an expected SEC peak and a matching CD trace for secondary structure. The five successful designs include A_1_1, A_2_1, A_2_2, A_3_1 and A_3_2; the remaining 19 designs are classified as failures. ESMFold in silico 'melting' curves constructed by masking the sequence to the language model from 30%-99%. The sigmoidal inflection point values (representing 'melting temperatures') of the melting curves were obtained.

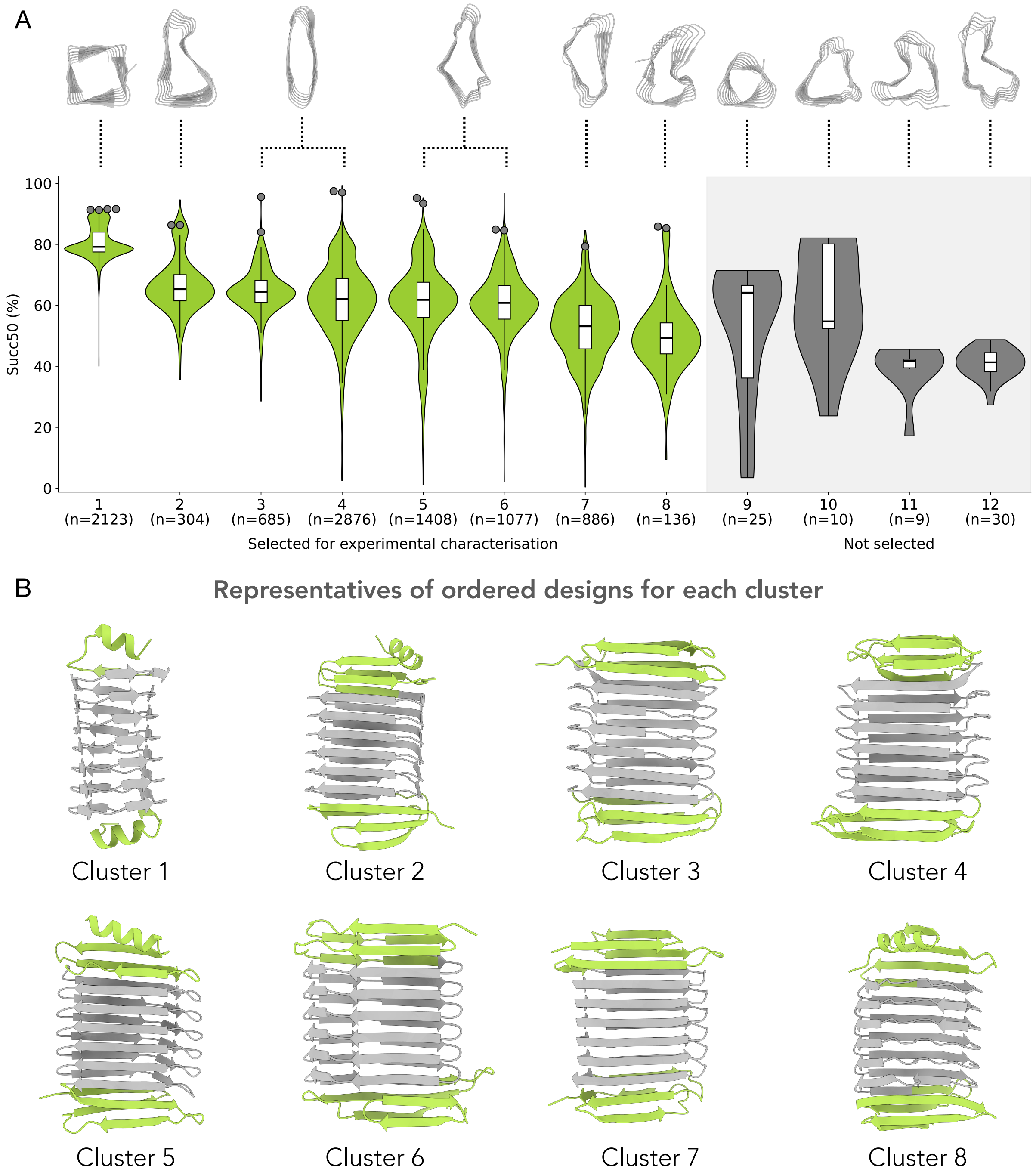


**Supplementary Figure 5 | *in silico* melting temperature of capped β-solenoid selection for experimental characterisation. (A)** Distribution of "melting temperatures" from ESMfold masking experiments for the capped β-solenoid designed clusters that passed previous quality metrics. The 12 clusters were made with Foldseek using a TM-score threshold of 0.7. The β-solenoid backbone from the in silico evolution loop that represents each cluster is shown above the violin/boxplot. The clusters are ordered based on median Succ50 (%) values. Sample sizes (n) for each cluster are indicated below each plot. The plots in green were the clusters that were used for experimental characterisation, while the grey plots were not carried forward for low successful sample numbers and lower meting temperatures. The top 4 sequences were taken from cluster 1, the top 2 were taken from cluster 2, 3, 4, 5, 6, 8 and the top one from cluster 7. **(B)** Best quality representative ordered from each cluster, caps are shown in green and solenoid region in grey.

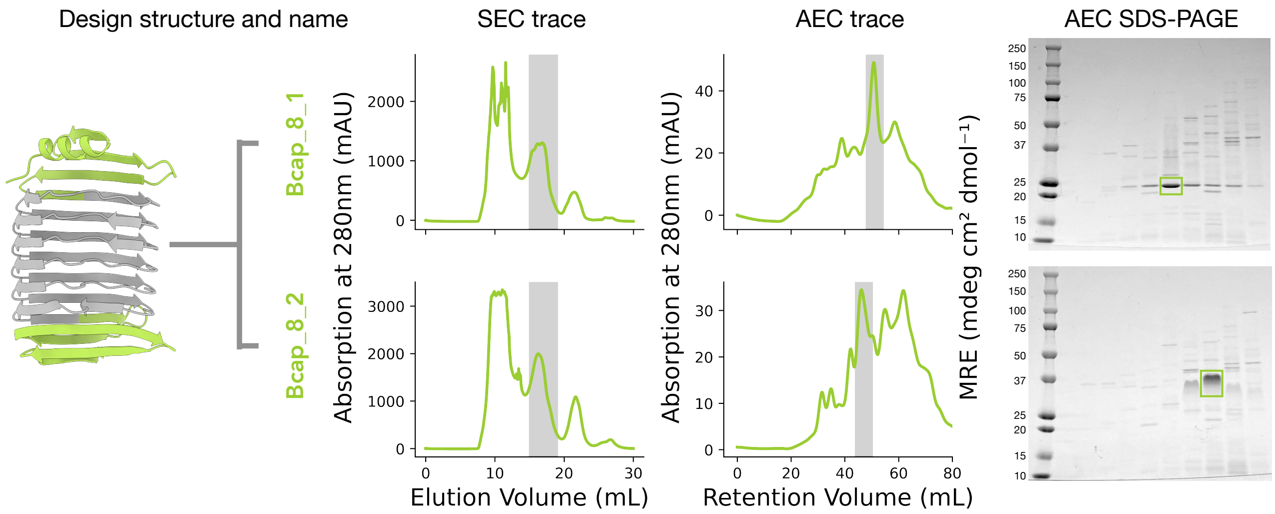


**Supplementary Figure 6 | Biophysical characterisation of capped β-solenoid proteins.** First panel, computational structure and name of the design characterised. β-solenoid region in grey and capping region in green. Second panel, SEC chromatogram. The highlighted region indicate the fractions pooled to for AEC. Third panel, AEC chromatogram, showing the elution profile of the pooled SEC fractions. This step is used to further purify the protein based on charge, with distinct peaks indicating elution of different species or forms of the protein. The highlighted region indicate the fractions pooled to continue for further characterisation. Fourth panel, SDS-PAGE of AEC fractions. Green boxes mark the band and fraction selected for further characterisation.

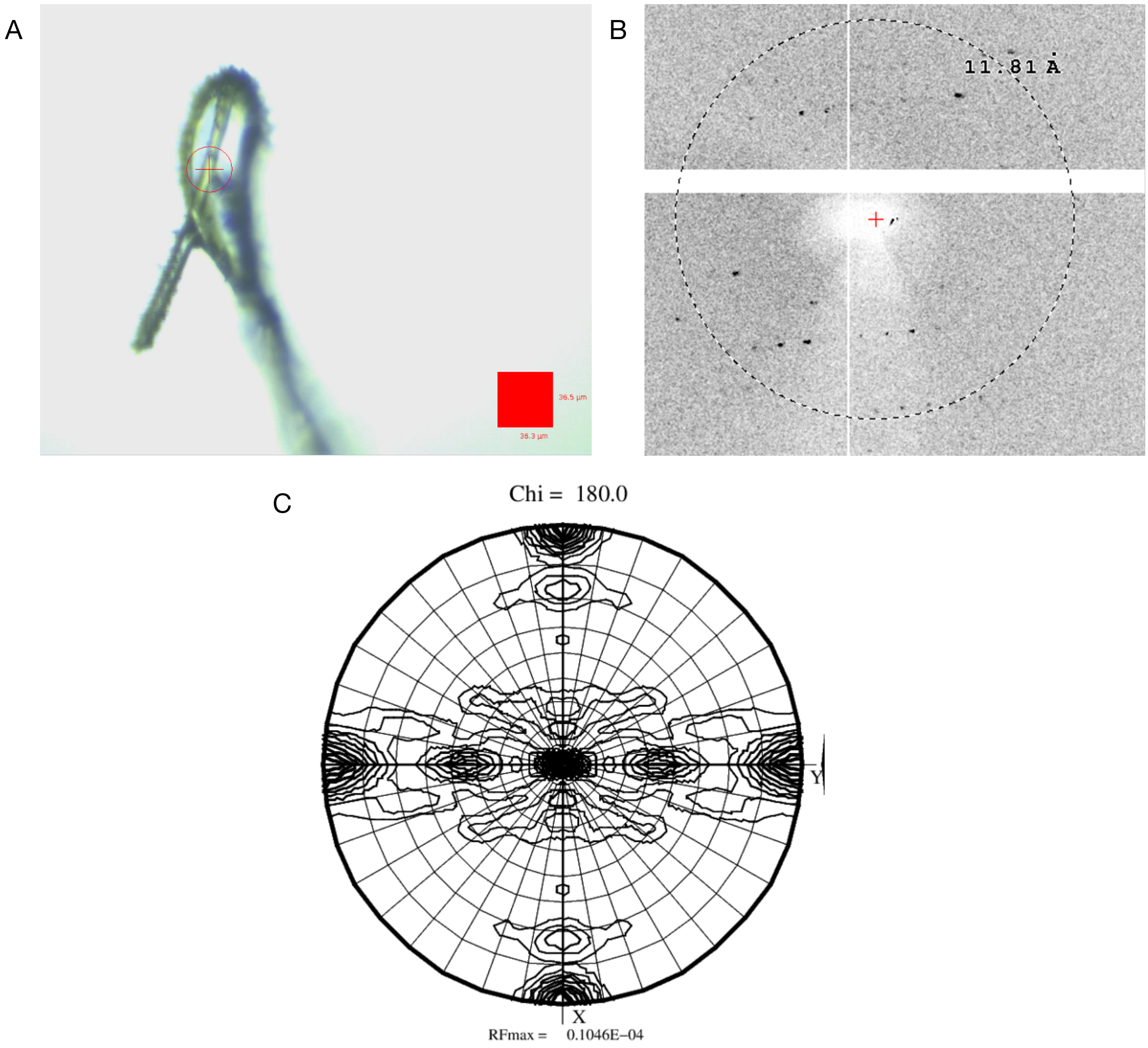


**Supplementary Figure 7 | Crystals and diffraction data from Bcap_8_2. (A)** Needle-like crystal mounted on the diffractometer on I24. Crystal is ~200 µm long and ~10 µm thick. **(B)** Detail of diffraction pattern from Bcap_8_2 crystal. **(C)** The unit cell parameters were a = 64 Å, b = 118 Å, c = 284 Å; α = β = γ = 90°, with an orthorhombic space group. The asymmetric unit was estimated to contain ~8–12 copies of the molecule. Analysis of the native Patterson function in Phaser showed no significant off-origin peak, suggesting an absence of major translational NCS. Molecular replacement attempts, with the design model of Bcap_8_2, did not yield a solution in any of the eight possible space groups. Diffraction data collected from Bcap_8_2 crystal showed weak features in the self-rotation function at ~10 Å resolution, suggesting the presence of at least two non-crystallographic 2-fold axes.

### Supplementary Tables

| Supplementary Table 1: DNA sequences of designed solenoid proteins from *in silico* evolution platform that were experimentally characterised | |
|---|---|
| **Design name** | **DNA sequence** |
| **AB_1_1** | ATGATGGGCAGCAGCCACCATCACCACCACCACTCCAGCGGTCTTGTGCCTCG<br>GGGTAGCagtTCAAATATTAAGACACTTACTTTTAACGTTGATACCCTCACACTGAC<br>GGTTGAGGAAGCGGAGAAGTACAGCAACGTCAAAACCTTGACATTTAATTCGCCA<br>ACCCTAACGCTGACAGTGGATATCGCGAAGGCGTTCTCTAATGTAGAAACCATTAC<br>CTTTAACACGCCCACCCTTACGCTTACCACGGATATTGCAAAGGCATTCTCAAACG<br>TGAAAACGCTCACCTTCAACACCCCAACTCTGACCTTAACCACCGAGATTGCCAA<br>GGCCTGGAGCAATGTGGAAACGATTACGTTTAATACCCCGACGCTAACTATTACTA<br>CCGAACAGGCAAAAGCTTGGTCGAATGTGAAGACCCTGATTTTCAATACGCCGAC<br>TTTGACCCTGTCGGATGAAGTCAAGAAAGCGCTGTAA |
| **AB_1_2** | ATGATGGGCAGCAGCCACCATCACCACCACCACTCCAGCGGTCTTGTGCCTCG<br>GGGTAGCagtGCGTCTATTGAAGAGTTGACCTTTAATGTGTCTAATTTGACGCTCAC<br>CGTAGAGGAAGCGAAGGCGTATAGCAATGTTAAGAAGCTTACTTTCAACACCCCG<br>AACCTGACCCTGTCTGTCGAAGTGGCCAAAGCGTGGAAGAACGTGAAGGAACTT<br>ACATTTAATACTCCAAACCTCACAATAACTACCGAAATCGCAAAGGCCTGGAAGAA<br>CGTAAAGAAACTGACGTTCAACACGCCGAATCTGACCATTACCACCGAGATCGCC<br>AAGGCTTGGAAGAATGTGGAGGAATTAACGTTTAATACCCCTAACCTTACTATCAC<br>CACTGAAATTGCAAAAGCATGGAAGAATGTCAAGAAAATCACTTTTAACACTCCCA<br>ACTTAACTTTATCCGAGGAGGTGAAGAAAGCTCTGTAA |
| **AB_2_1** | ATGATGGGCAGCAGCCACCATCACCACCACCACTCCAGCGGTCTTGTGCCTCG<br>GGGTAGCagtATGAAATCTATTACACTGAAGAATATCGGCAAATTCCCACCGGAGTC<br>GTTCCTAGAACTGTTTAAGAACCTCGAAGAGGTGACACTTATCAACCCGGACAAA<br>TGGCCACCCGAAACGTGGGTGCCGTTGTTCGAGAAGCTCAAGAAAGTGACCTTA<br>GTGAACATCGATGCCTGGCCGCCTGAAACCTGGGTACCGCTGTTTGAGAAATTG<br>GAGGAAGTCACGTTGGTAAATATTGACGCGTGGCCGCCGGAAACTTGGGTCCCG<br>CTCTTTGAAAAGCTGAAAGAAGTTACCCTCGTTAACATTGGTGCATGGCCTCCTG<br>AGACATGGGTTCCGTTATTTGAAGGCTTAGAAAAGGTTACTATTGTGAATCCTGGT<br>CCTTGGCCGCCAGAGAGCTGGGCGGTTTTATTCGAAAAGTAA |
| **AB_2_2** | ATGATGGGCAGCAGCCACCATCACCACCACCACTCCAGCGGTCTTGTGCCTCG<br>GGGTAGCagtATGTCGTCCATCGAATTGAAGAATATCGGTAAATATAAGCCAGAGTC<br>ATTTCTGGAGCTATTCAAAGATCTGAAGAAAGTGACGCTGATCAACATAAACAAGT<br>GGCCACCGGAGTCCTGGATACCGTTGTTCGAAAATTTGGAGGAAGTTACCTTAGA<br>AAACATCAATAAATGGCCGCCTGAAACATGGATCCCGTTATTTGAGAATTTAAAAGA<br>GGTGACACTCAAGAACATTAACAAATGGCCGCCGGAAACGTGGATTCCGCTGTTT<br>AAGAACCTAGAGGAGGTCACGCTCGAGAACATTGGCAAATGGCCTCCTGAGACT<br>TGGATTCCTATTTTCAAGGGATTAAAGAAGGTTACTATTATTAACCCTGGTCCTTGG<br>CCCGAAGAGAGCTGGAAGATTCTGTATGAACAGTAA |
| **AB_3_1** | ATGATGGGCAGCAGCCACCATCACCACCACCACTCCAGCGGTCTTGTGCCTCG<br>GGGTAGCagtAGCGCTAAATATAAGGGCAAAACGATTTCTAACGACGACAACTTCG<br>CATTGAATCCAAAGAAGTATCCGCTGGAAGATCTGATTGGTCTGACCTTTAAGAATA<br>TCCCGAACCTCGCCAATAACCCGAGCAAGTGGCCCACCGAGGGCCTGAAGACG<br>CTGACAATTATAAACAACGAAAACTTTGCGAACAATCCGTCGAAATGGCCGACTGA<br>GGGTTGGAAGGGACTCACCATCGAGAACAACAAGAACTTTCCGAATAATCCCTCT<br>AAGTGGCCTACGGAACAGTGGAAAGGCTTAACCATAAAGAATAATGAGAATTTTGC<br>TAAGAACCCAAGTAAATGGAAAGCTGAGCAGTGGAACGGCATTACCCTGGAGAA<br>CATTCCTAACTACAATAACAACCTTAATACCTGGCCGTAA |

| Supplementary Table 1: DNA sequences of designed solenoid proteins from *in silico* evolution platform that were experimentally characterised | |
|---|---|
| **Design name** | **DNA sequence** |
| **AB_3_2** | ATGATGGGCAGCAGCCACCATCACCACCACCACTCCAGCGGTCTTGTGCCTCG<br>GGGTAGCagtGCGGCTACTATGGTCGGTAAAACCATCTCTAATGATCAGAACTTCG<br>CGTTAAACCCGCAGGATTACCCACTCGAGGATCTCAAAGGCTTGACTTTTAAGAA<br>CCAAAACAACTTGGCGGACAATCCGTCTCTTTGGCCACTTGAAGGCCTGAAAAC<br>GCTGACGATCAAGAATCAACCTAATTTTGCAGATAACCCTAGCCTCTGGCCCCTG<br>GAGGGATGGAAAGGGCTCACGATCGAAAATCAGCCTAACTTTGCTAAGAACCCC<br>AGTCTGTGGCCGTTAGAACAGTGGAAAGGTCTCACCATTAAGAACCAACCGAATT<br>TCGCTAAATATCCAAGCTTATGGAAGAAGGAACAATTGGAAGGGGTGACTATTGTG<br>AACCAGCCGAACCTGGCGAAATACCCGGATAAATGGCCGTAA |
| **A_1_1** | ATGATGGGCAGCAGCCACCATCACCACCACCACTCCAGCGGTCTTGTGCCTCG<br>GGGTAGCagtAGCTGGGAAGAGGAGTTGGAAAAGTTTATTCGCTATCTGAAAGAGA<br>ACGATCTGCCCGAGGAGGAAAAGGAGAAGCTTGTGCGCGAGTTTCTGCGCAAA<br>GCTAAGGAAAATTTGAACCCGGAGGAATTATTTAAAGTGTTTCTACGTATCATCAAT<br>GAAGTGCCGTTGCCAGAAGAATTCAAGAAGAAATTGGTGGAAGAGTATTTTGAAT<br>GGGCGAAAGAAAACCTCAACCCCGAAGAACTGTTTAAGGTGTTCCTGCGTATTAT<br>CAACGAAGTACCATTACCAGAGGAATTTAAGAAGAAGTTAGTTGAAGAATATTTCGA<br>GTGGGCCAAGAAGAACCTGAACTTGGATGAGCTGAAGAAAGTCTTTGAGAAAAT<br>CCTAAAAGAGGTGCCTATACCGGAAGAGTTCAAGGAGGAGCTTAAGAAACGTTAT<br>GAGGAATACATGAAGAAGCGCCAGTAA |
| **A_1_2** | ATGATGGGCAGCAGCCACCATCACCACCACCACTCCAGCGGTCTTGTGCCTCG<br>GGGTAGCagtAAGAGCGAAGAGGAGCTCGAAAAGTTCATTAAGTACATCAAAGAAA<br>ATAACCTACCCGAAGAGGAGAAAGAAAAGCTTGTAGAAGAATTTCTGGAGTACGC<br>CAAGAAGAATCTAAACCCGGAAGAACTGTTCAAAGCGTTCCTGAAAATTATAAAGG<br>AGATCGACCTGCCAGAGGAAGTGAAGGAGAAACTGGTGGAAAAGTTCTTTGAGT<br>GGGCCAAGAAGAACCTCAACCCCGAGGAACTATTTAAAGCCTTCCTTAAGATTATT<br>AAAGAGATTGACTTACCGGAGGAATTCAAGGAGAAGCTCGTCGAGAAATTCTTCG<br>AATGGGCTAAGAAGAATCTGAACCTGGAAGAGTTGAAGAAGTATTTCAAGAAGATC<br>CTAGAGGAAATCGACTTGCCTGAGGAGTTTAAGAAGAAGTTAAAAGAGAAATATGA<br>GGAATATATCAAGAAGAAGTCCTAA |
| **A_2_1** | ATGATGGGCAGCAGCCACCATCACCACCACCACTCCAGCGGTCTTGTGCCTCG<br>GGGTAGCagtCCGGTGCCGCCGGAATTGCGGCCAGAACTAGAGCGCGAAGAGAT<br>CCTCGAGTGGATTAAACAGGGTAAGGAGTTTGACTGGGAGGCACGGTTAGAAGA<br>AGTATTCCCGGAGGAAGAACGGCCCGAAGCGATGCTTAAAATTATACTGTTATGGA<br>TCGAGCAGGGGAAAGAATTTGATTACGAGGCGTTACTGGAGAAATGGTTCCCTGA<br>GGAGAAACGTCCGGAAGCCATGCTCAAGATCATATTGCTGTGGATAGAACAAGGT<br>AAAGAATTCGATTATTTTGCGCTGCTTGAAAAGTGGTTTCCAGAAGAAAAGCGTGA<br>AGAGGCTAAGGAAAAGATTCTCGAACTGATGAAGAAGCAAGGCAAAGAGGTGGA<br>CTTAGAGGAATGGGAGAAGCGCTAA |
| **A_2_2** | ATGATGGGCAGCAGCCACCATCACCACCACCACTCCAGCGGTCTTGTGCCTCG<br>GGGTAGCagtAAAATTCCTAAGGAGAAAATACCGGAACTCAAAGAAAAGGAAATCC<br>TGGAGAAGATCAAGAATAACGAAAAGTTTGATTATGAAAAGATTATCGAGAAGACCT<br>TTCCCAAAGAGAAGCGCCCCGAAGTGATGTTAAAGATAATAAAGCTCATGATAGAA<br>CAGAACGAGAAATTTGACATCGAAAAGCTTATTGAGAAGTTCTTCCCCAAGGAAAA<br>GCGGCCAGAGGCCATGCTGAAAATCATTAAACTGATGATCGAGCAGAATGAGAAG<br>TTCGACATAGAGAAGCTGCTGGAAAAGTTCTTTCCGAAAGAGAAACGCGAGGAG<br>GCTAAGAAGGAGATCCTAGAATTGATGAAGAAGAAGAAAATTAAGGTGGACCTGA<br>AGAAATGGGAGAGCCGCTAA |

**Supplementary Table 1: DNA sequences of designed solenoid proteins from *in silico* evolution platform that were experimentally characterised**

| Design name | DNA sequence |
|---|---|
| **A_3_1** | ATGATGGGCAGCAGCCACCATCACCACCACCACTCCAGCGGTCTTGTGCCTCG<br>GGGTAGCagtTCAAGCATCCCGCTGAGTGAAATCAGCGAAGAAGAGTTTCTGAAA<br>ATCCTGGATGAGATGGAAGAAAACGGTAAGTCCAAGGAGGAGATCCTAAAGAAAG<br>CGTTAGAGTACATAGAGAAGAAAGACCCACAGACCATCTCGGACGAATTGTTCAA<br>CCGCTACATTAACCTGATGAAAGAGCTGGGCAAAAGCAAAGAAGAGATTCTCAAC<br>GAAATGGAGAAATATATCGACAAGAAAGATCCGCAAACGATCAGTGACGAGTTATT<br>CAATTACTACTTAAATCTGCTGAAGGAGCTTGGCAAGTCGAAAGAGGAAATTTTAA<br>AGGAACTGAAGAAATACATTGATAAGAAGGACCTTAAGACTATCTCCGACGAACTC<br>AAGGAGTATTACATCAACTTATTAAAAGAACTGGGGAAAAGTGAGGAGACAATCGA<br>GAAGTATAAGAAGAAGTTCGAGTAA |
| **A_3_2** | ATGATGGGCAGCAGCCACCATCACCACCACCACTCCAGCGGTCTTGTGCCTCG<br>GGGTAGCagtGGCAGCATCCCGCTGTCGCTGATTAGCAATGAGGAGTTTATGAAAA<br>TCCTGGAGGAAATGAAGAAGGAAGGCAAAAGCCTCAAGGAGATATTGAACGAATT<br>GGAAAAGTGGTTGGAGTCCAAAGATCCGCAAACCGTGTCCGATGAGTTATTCGA<br>GTACTATCTAAATTTGATGAAGGAACTCGGCAAGTCCTTAGAAGAAATTCTCAAGAA<br>GCTTAAAGAATGGTTAGAGAAGAAGGATCCACAGACAGTCTCCGACGAACTGTTT<br>GAATATTATCTGAACCTCATGAAAGAGCTGGGTAAGAGTCTGGAAGAGATTCTTAA<br>CTTTGCACTAGAGTGGCTCGCGAAGAAAGACCCGCAGACCGTTAGCCCGTCGTT<br>GGCGGAGTATGTGCTCAACTTAATGAAACAATTAGGTAAATCGCAAGATACCATCG<br>ACGCGCTAAAGGCGGCGCTGGCGTAA |
| **B_1_1** | ATGATGGGCAGCAGCCACCATCACCACCACCACTCCAGCGGTCTTGTGCCTCG<br>GGGTAGCagtAGTAAAATCGAAGCGGAGGAGATCGAGGCAGAAGAAATAAAGGCT<br>AAGGAAATCAAGGCGAAAGAGATTAAAGCGAAGAAATTTAAAGCAGACACTATAGA<br>GGCGGATACTATTAAAGCGGATGAAATTAAGGCGGATAAGTTTGAAGCCGATACATT<br>TAAAGCAAAGACGATAAAAGCCAAGAAATTCAAGGCAAAGAAGTTCGAAGCAGAT<br>ACCATAGAAGCCGACACCATTGAGGCGGACGAAATTGAAGCGGACAAATTTGAA<br>GCTGACACATTCGAGGCCAAAACCATCAAAGCTAAGAAATTTAAGGCCGACAAGT<br>TCGAAGCGGACACGTTCAAAGCTGATACGTTTGAGGCGAAGGAGTTCAAAGCTA<br>AAGAATTCGAGGCTGAGACTTTTGAAGCCGAAAAGTTTAACGCGTAA |
| **B_1_2** | ATGATGGGCAGCAGCCACCATCACCACCACCACTCCAGCGGTCTTGTGCCTCG<br>GGGTAGCagtGCCGAAATAAAAGCAGAAAAGTTCGAGGCGGAGGAGTTCAAGGC<br>CAAAACATTCGAAGCGGATAAGTTTGAGGCCGATGTGTTTGAAGCCAAGACTTTC<br>AAAGCCGATACCTTTAAGGCGGGCGAGTTCAAGGCTAAGAAATTCAAAGCTAAAA<br>CGTTTAAGGCAGGGACCTTCGAAGCAAAGAAGTTTAAAGCTGAGGAATTCGAAGC<br>CGACACTTTTGAAGCCGGCACGTTCAAAGCGGGAGAATTTGAGGCAGATAAATTT<br>AAGGCGAAGACATTTAAGGCTGGCACTTTCGAAGCAGGCAAAATTAAAGCGAAGA<br>AGATAGAAGCTAAGACTATTAAGGCCGGAACCATTACCGCGAAAACCTTTGATGCG<br>GACGAATTCACCGCAGACAAGTTTGAGGCTGATACGGTTAGCGCGTAA |
| **B_1_3** | ATGATGGGCAGCAGCCACCATCACCACCACCACTCCAGCGGTCTTGTGCCTCG<br>GGGTAGCagtGCGAGTTTCACTGCCGATAAGTTCGAGGCAGAAGAATTTGTCGCC<br>GAGACGTTCAGCGCAGACACCTTCACAGCACAAACATTCAAGGCCAAAACGTTT<br>AAGGCCAACACTTTCACTGCTGGTACCTTTACCGCCGACACATTTGAAGCTGACA<br>CCTTTGAGGCGAACACGTTTACCGCGGGCACATTTACCGCGAAAACCTTCAAAG<br>CCAAGACCTTTACAGCGGATACATTCACCGCTGGCACCTTTACGGCTGATACGTT<br>TGAAGCAGGCACTTTTAAAGCAAATACCTTTACAGCTGGGACGTTCACGGCAAAG<br>ACATTTAAAGCGAAGACTTTCGAAGCTAAAACTTTTACGGCCGATACTTTTGATGCT<br>GATACCTTTAGTGCAGATACCTTTACTGCGGACACGGTGGAAGTGTAA |

| Supplementary Table 1: DNA sequences of designed solenoid proteins from *in silico* evolution platform that were experimentally characterised | |
|---|---|
| **Design name** | **DNA sequence** |
| **B_2_1** | ATGATGGGCAGCAGCCACCATCACCACCACCACTCCAGCGGTCTTGTGCCTCG<br>GGGTAGCagtGCGGAATATAGCTACAATGATAACTCGACCCGCACCACCGCGGAG<br>ACCATTACGTACACAGCGCCGGAGGAAGCGAAGAATTTCACGATCAACAACAACT<br>TCACCTCCACTGCCCCGTTTAATCTGACCTTCAACGCGCCAAAGAAAGCGGATAA<br>CTTTACGGTGAACGACAACTTTACAACCACGGCGTCATTTAACATCACGTATAACG<br>CCCCAGAAACAGCTAAGAACTTCAATATCAATAATAACTACACCTCTAAAGCCTCAT<br>TCAACATAAACTATAATGCGCCCAAAACCGCAGACAATTTTAACATTAACGATAATTT<br>TACCACTACGGCCTCTTTCAACCTGACATATACCGGTCCAGAGACAGCCAAGAAC<br>TTCTCAGTTAACCTCAACTCTACGACGACCGCCAGCAGCACTGTGACTATTACTG<br>AACCGAAAACTGTGTAA |
| **B_2_2** | ATGATGGGCAGCAGCCACCATCACCACCACCACTCCAGCGGTCTTGTGCCTCG<br>GGGTAGCagtAGCACCTACACCACGAACGACAACAGCACGCTTACCACTACTAAG<br>AACATTACGTATACAGCGCCGGCCACCGCGACCACATATACCGTGAACTTAAACAT<br>GACCCTCACGGCTCCCTTCACTCTGACTGTCAACGGCCCAACGACCGCAACAA<br>ACTTTACCGTAAATGATAACATGACAATCACAGCCTCTTTCACCCTGACCTATAACG<br>CCCCGACGACGGCAACCAATTATACGATAAACCTCAATATGACGAGCACTGCGAG<br>CTTTACATTGACCATTAACGGGCCAACTACGGCGACTAATTTTACTGTGAATATCAA<br>CTCCACCTTAACTGCCTCGTTTACCCTAACCATCACCGGTCCGGAAACCGCTACG<br>ACTTACACTGTTAACCTGGATTCGACCCGTACTGCAAATTCTACCATAACCTTGAC<br>GTCTCCGGCTACAGTGTAA |
| **B_2_3** | ATGATGGGCAGCAGCCACCATCACCACCACCACTCCAGCGGTCTTGTGCCTCG<br>GGGTAGCagtTCCACGTATGAAACCAACGATAACAGCACGACCACCACGGCCCAA<br>ACAATTACCTATAGCGCGCCAGAGACGGCCGATACGGCGACGATTAACGACAACT<br>CTACGTCCACTACTAGCTTCACCCTGACATATAACGCACCGAAAACCGCGAAGAA<br>CTTCACCATCAATGATAAGACGAAAAGCACTGCGAGCTGGACAAAGACCTACAAC<br>GCCCCAGAAACAGCGGACACCGCCACGATCAACGATGATTTCACGTCTACCGCA<br>AGCTTTACCAAAACATACAATGCGCCTAAAACGGCTAAGAATTTTACAATAAATATTA<br>ATGCCAAATCGACCGCTTCTTGGACCTTGACCATAACTGCCCCGGAGACAGCTG<br>ATACTTATACCGTCAACCTGAATACCACTTCGACTGCGGATAGTACCCTCACCGTT<br>ACTGCACCTAGCACAGTGTAA |
| **B_3_1** | ATGATGGGCAGCAGCCACCATCACCACCACCACTCCAGCGGTCTTGTGCCTCG<br>GGGTAGCagtGAAATTGAGTTTGAAAACCCCAAGGAGTTCGTGCTTTATAAGGATTA<br>TACGTTCGACGTGGATGAATTCACGCTGAAAGGGAAATTTAAAGAGCCCGAGAAA<br>TTTTACGACTTTGAGAACTCGACATTTAAAGTTGACAAGGTCACATATGAGTATACA<br>TTCGAAAAGCCGAAGAAGTATTACCTGTTCAAGAACAGTACCTTCGACGCCAAGG<br>AAGTCATTATTAAGGGGAAGGGCAAAGAACCGGAAGAGTTCTATGATTTCGAAAAC<br>AGCACGTTTAACGTTGATAAGGTTGAATACAAAGGCGAATTCGAGAAGCCTAAGAA<br>GTACTATATTTTCAAGAATAGCACTTTTAATGCGAAGGAGGTGATCGTAGACCTGAA<br>GATCGAGAAGCCGGAGAAGTTTTATTTGTTCGACAACTCCACCTTCAATGTGGAC<br>AGCATCAAACTGTAA |
| **B_3_2** | ATGATGGGCAGCAGCCACCATCACCACCACCACTCCAGCGGTCTTGTGCCTCG<br>GGGTAGCagtGATATCGAGTTTGAGAACCCCAAGAAACTGACCCTGTACAAGGAC<br>AAGACCTTCAATGTAGAAGAGGTCAAATTAAACCTGAAATTTAAGAAACCGGAAGA<br>GTTGTATTTGTTCGATAATTCGACGTTTAACGCGAAGAAAGTCGAGATTAACATTAA<br>GTTCGAGGAACCAAAGAAATTGTACTTCTTCAAGAATAGCACCTTTAACGCAGACA<br>AAGTCATTTTCAAGCTGGAGGGCAAGAAGCCGGAGGAGCTGTATCTCTTCGACA<br>ACAGTACGTTCAACGCTAAGGAAGTGAAATATGAAGCGAAATTCGAAGAACCTAA<br>GAAGCTTTACGTGTTTAAGAACAGCACATTCAATGCGGATAAAGTAATCGTGGATC<br>TGAAGATCAAGAAGCCTGAGGAACTCAAGCTCTTTGATAACACCACTTTTAACGTG<br>AAGGAGATTGAGATCTAA |

**Supplementary Table 1: DNA sequences of designed solenoid proteins from *in silico* evolution platform that were experimentally characterised**

| Design name | DNA sequence |
|---|---|
| **B_3_3** | ATGATGGGCAGCAGCCACCATCACCACCACCACTCCAGCGGTCTTGTGCCTCG<br>GGGTAGCagtGAGCTCGAGTTTGAAAACCCAGAAGAATTCACCCTGTACAAGAATA<br>AGACATTCAACGTCGATTCGGTGACCTACAAAGGGAAGTTCAAGAAACCAAAGAA<br>ATTCTATCTATTCGAGAACAGCACATTTAACGCGAAATCTGTGACGTACGAATTTGA<br>AGGCGAGGAACCGGAAGAGTTCTACCTCTTTGACAACAGTACTTTTAATGTAGATT<br>CTGTTACTTATAAAGGCAAATTTAAGAAGCCCAAGAAGTTTTATTTGTTTAAGAACTC<br>CACTTTCAATGCGAAGTCCGTTACGGTGGAACTGGAGATGGAAAAGCCAGAGGA<br>ATTTTACCTGTTTGAGAATAGCACCTTTAATGCCGATAGCGTGACTATCAACCTGAA<br>GATCGAAGAACCTAAGAAACTGAGCTTGTTCGAAAACACTACGTTTAACGTGAAA<br>CGGGTCGAGATTTAA |
| **B_4_1** | ATGATGGGCAGCAGCCACCATCACCACCACCACTCCAGCGGTCTTGTGCCTCG<br>GGGTAGCagtACTATCACCGGCGGTACGATTACCGGAGGTACCATTTCGGGAGGG<br>ACCATCACGGGCGGGACAATCATCGGCGGTACTATCATAGGTGGCACGATAAAAG<br>GCGGAACCATAATCGGTGGCACGATCATTGGCGGGACTATAGAGGGCGGCACCA<br>TTATAGGCGGAACAATAATTGGCGGCACCATCAAAGGTGGGACGATTATAGGAGG<br>CACCATCATTGGCGGCACGATTGAAGGCGGAACGATCAATGGAGGCACGATTATT<br>GGTGGTACAATTAAGGGTGGCACAATCAACGGCGGTACCATTATCGGAGGAACTA<br>TCGAAGGAGGTACCATAAACGGAGGCACTATTATTGGAGGTACGATTAACGGTGG<br>CACTATTAATGGCGGCACCCGCAGCGGTGGAACCCTGATTCCGGATTAA |
| **B_4_2** | ATGATGGGCAGCAGCCACCATCACCACCACCACTCCAGCGGTCTTGTGCCTCG<br>GGGTAGCagtACCATAACGGGAGGAACAATCATCGGCGGGACGATCACTGGAGGT<br>ACTATCAGCGGTGGCGTGATCATAGGTGGGACCATCACCGGAGGCACCATTTCG<br>GGTGGTGTCATCATTGGCGGCACGATAACCGGTGGCACAATTTCAGGAGGCGTTA<br>TAATTGGTGGCACCATTACCGGCGGGACAATAATCGGCGGCGTCATAATAGGCGG<br>AACGATCACGGGCGGCACTATCATTGGTGGGGTCATTATAGGAGGGACTATCACA<br>GGTGGCACCATCATTGGAGGTGTTATTATCGGAGGCACTATTACAGGCGGTACCAT<br>TATTGGCGGAACTATAATTGGCGGTACGATTACTGGTGGAACCATTATCGGTGGCA<br>CGATTATTGGTGGTACAATTACGGGTGGCACTGTTACCCCGGATTAA |
| **B_4_3** | ATGATGGGCAGCAGCCACCATCACCACCACCACTCCAGCGGTCTTGTGCCTCG<br>GGGTAGCagtACTATCACGGGCGGTACCATAACCGGTGGCACAATCACTGGTGGC<br>ACGATTTCCGGCGGGACCATCATCGGTGGTACGATCTCTGGCGGAACGATATCC<br>GGAGGAACAATCATAGGCGGGACGATCATAGGAGGAACTATTTCGAACGGGACCA<br>TAATTGGAGGGACAATAATCGGTGGCACCATCTCAGGCGGCACCATTATAGGTGG<br>AACCATCATTGGCGGCACTATAAACGGAGGTACAATTATTGGCGGCACCATCAAG<br>GGAGGCACGATCAACAACGGCACTATCATCGGAGGCACCATTGAAGGCGGCAC<br>GATTAACGGTGGTACTATTATCGGCGGCACTATCAAAGGTGGGACTATTAATGGCG<br>GTACCATTATTGGTGGTACCCGCGAGGGTGGCACGCTGAATCCAGATTAA |

| Supplementary Table 2 \| Data collection and refinement statistics for A_1_1 | |
|---|---|
| | **A_1_1** |
| **Wavelength** | 0.62 |
| **Resolution range** | 27.96 - 2.833 (27.96 - 2.83) |
| **Space group** | I 41 |
| **Unit cell** | 74.841 74.841 50.878 90 90 90 |
| **Total reflections** | 21865 (21865) |
| **Unique reflections** | 3395 (3395) |
| **Multiplicity** | 6.4 (6.4) |
| **Completeness (%)** | 98.95 (98.95) |
| **Mean I/sigma(I)** | 6.08 (6.08) |
| **Wilson B-factor** | 81.74 |
| **R-merge** | 0.1729 (0.1729) |
| **R-meas** | 0.1885 (0.1885) |
| **R-pim** | 0.07399 (0.07399) |
| **CC1/2** | 0.989 (0.989) |
| **CC*** | 0.997 (0.997) |
| **Reflections used in refinement** | 3377 (3377) |
| **Reflections used for R-free** | 159 (159) |
| **R-work** | 0.2570 (0.2570) |
| **R-free** | 0.3000 (0.3000) |
| **Number of non-hydrogen atoms** | 1259 |
| **macromolecules** | 1259 |
| **ligands** | 0 |
| **solvent** | 0 |
| **Protein residues** | 151 |
| **RMS(bonds)** | 0.002 |
| **RMS(angles)** | 0.46 |
| **Ramachandran favored (%)** | 96.64 |
| **Ramachandran allowed (%)** | 3.36 |
| **Ramachandran outliers (%)** | 0.00 |
| **Rotamer outliers (%)** | 0.00 |
| **Clashscore** | 2.81 |
| **Average B-factor** | 103.09 |
| **macromolecules** | 103.09 |

*Statistics for the highest-resolution shell are shown in parentheses.*

| Supplementary Table 3: DNA sequences of designed β-solenoid solenoid proteins with caps that were experimentally characterised | |
|---|---|
| **Design name** | **DNA sequence** |
| **Bcap_1_1** | ATGATGGGCAGCAGCCACCATCACCACCACCACTCCAGCGGTCTTGTGCCTCG<br>GGGTAGCagtACAACCCTGACCGCGGCGGAGCTGGAAGCGATTGCAGCCAAGG<br>GCGGTACAGTGAAGAATGCGACAGTAGAAAATCAGACCTTCAAGAACCAGACGTT<br>TGATAATGTTACATTCGAGCAGGTGACATTTAAGAACGTAACCTTCCGGTCGGTGA<br>CGTTTCGCAACGTTACTTTCAAGAACGTGACATTTGAATCCGTGACGTTCGATAGC<br>GTGACCTTCGAAAATGTGACCTTTAAGAACGTGACGTTTGAAAACGTGACGTTCA<br>AGAACGTAACTTTTGAGCAAGTGACCTTTGAGAATGTGACGTTTAAACAGGTTACC<br>TTTGATCAGGTCACCTTTCGGAACGTGACTTTCCGCAATGTGACCTTCGACAATG<br>TGACTTTTAAAACCGTTACCTTTGACAACGTCACTTTTCAGAACGTGACCTATTCTA<br>ACGTTACCTACGAATCGGTTACTCTAAAATCTGTGACTTACCGGCCTGGTAATACAT<br>ATAACAACTTAAGCGGCACCATAACCATTACGGGCACGATTACTTATACCtaa |
| **Bcap_1_2** | ATGATGGGCAGCAGCCACCATCACCACCACCACTCCAGCGGTCTTGTGCCTCG<br>GGGTAGCagtACCACCCTTACCGCGGCGGAGTTAGAAGCGATTGCAGCCAAGGG<br>CGGCACGGTAAAGAATGCGACCGTAGATAATCAGACTTTCAGAAACCAGACGTTT<br>GAATCCGTGACCTTTGAGAACGTTACATTTCGTCAGGTTACCTTCAAGAACGTAAC<br>CTTTCGCACCGTCACGTTTAAGAATGTTACGTTCGATAGCGTGACATTTGATAACG<br>TGACATTCGAAAACGTGACGTTCAAGAACGTTACCTTTGAAACAGTCACTTTTAAA<br>ACCGTTACCTTCGAGAACGTGACGTTTGAAAATGTGACCTTTAAGCAGGTGACCT<br>TTGACAGCGTCACCTTCCGCAACGTGACATTTAAATCTGTTACTTTTGAGAATGTG<br>ACGTTCCGAAACGTGACCTTCGACAATGTAACTTTCCAGAACGTGACATATAACAC<br>TGTGACTTACGAGAGCGTTACCCTGAAGAACGTTACGTATAAACCCGGCAATACCT<br>ACAACAACCTCAGCGGGACCATTACTATTACGGGTACCATCACCTATACCtaa |
| **Bcap_1_3** | ATGATGGGCAGCAGCCACCATCACCACCACCACTCCAGCGGTCTTGTGCCTCG<br>GGGTAGCagtGCGACCCTGACAGTGACGGCCACATCGACTGGCAATGTGACTGT<br>AACCTATGATGCAGCCACGAGCACCCTAACTGTCACGATCAGCTCGGCCGCGGG<br>CCAGACGTTGAACGCGACGGTTACTATCTCTGCTACGGGTACGGTGACCTTATCT<br>AACATAAACATTAACATCACGTCAACTGCAAGCGGGACCGTGAACGTTAACATTAG<br>CGGAACGATTACTGCCGGAAACATAAACATCAACATTAACTCCACCGCCCCAATG<br>ACGATAAATATAACTTTCTCCGGTACCTTTACGGCGACTAATATAAACCTGAACATCA<br>ATAGCACGGCTCCTATGACTCTGAATATCACCTTCAAAGGGACATTCACAGCCGG<br>GAATATTAACATCAATATCAACAGCACTGCGCCGATGACAATTAACATTACGTTCGA<br>AGGCACGTTTACAGCTGGGAACATTAATATCAACATAAACTCGACCGCGCCCATGA<br>CCATCAACATTACTTTCAAGGGCACTTTTACCGCAGGCAACATCAACATCAATATTA<br>ATTCCACTGCTCCAGTTACACTTAACATTACATTTAAGGGTAACATAACAATTGGAAA<br>ACTGAAGATCGAATTCAAGAATGCGTCCGATTCTACCACGCTTACAATCACAACGG<br>AGTCTTCCGGGGTTACGACCACCACTACGTATACTGGTATTAAGAACGGCACCGTT<br>ATTGAGATTGATTCGCTCACTATTACCAACACCtaa |
| **Bcap_1_4** | ATGATGGGCAGCAGCCACCATCACCACCACCACTCCAGCGGTCTTGTGCCTCG<br>GGGTAGCagtGGTAAAGTGACGGTCGAGTCCAATGGCAAAACTATTACTATCGAGC<br>TGCCGGGCTATACGCTGACCAAACTGGAGTTAAATTCGAACGGTAAACTCGTGATT<br>ACCTTCAAAGACGAGAACGGAAACGAGAAAACCGTAGAGATCGACGGCAAGAAC<br>ATCACTAAAATCGAAATCAACCTGAAGGCGAAGGAGAACGTGTCGACTGAGATCA<br>AGCTGGAGAACATGAAGAACCTCGAGAAGATCAACATAAATATCGACGCCGAGAA<br>GAACCTGAAGCTTAACCTGACAATGAAGAATGTGAAGAACCTCAAGGAAATAAAC<br>ATCAATATTAACGCGAAAGAGAATTTGGAACTGAACATAGAGCTAGAAAACGTTAC<br>CAATCTAAAGAAGATCAACATCAACATAAACGCGGAGAAGAATCTGAAACTCAACC<br>TTAAGATGAAGAACGTGAAGAACTTAAAGGAGATTAATATCAATATAAATGCAGGTG<br>AGAACGCCGACATTAACATCGAGATGGAGAACGTCACCAACCTGGAAAGCATTAA<br>CATTAACATCAACGCTAAGAAGAACCTAAACATCAACATCAAGTTCAAGAACGTGG<br>ACGACTTCAACTTGAATATTAACATTACGACCAGCAACGGGAACACCATCACACTG<br>AACAAGAAAATAACCAAGAATATCACCGAAATCAAAATTAACGGCGAGGAAAAGAA<br>CGGCAATCTGACGCTCAACATTGAAATTACTTATAAAtaa |

**Supplementary Table 3: DNA sequences of designed β-solenoid solenoid proteins with caps that were experimentally characterised**

| Design name | DNA sequence |
|---|---|
| **Bcap_2_1** | ATGATGGGCAGCAGCCACCATCACCACCACCACTCCAGCGGTCTTGTGCCTCG<br>GGGTAGCagtATGAAACTAGAGGTCAAGGAGGAAGAGGGCGAACTGGAAATTAAA<br>ATATCGGGCGGGAACGTGACAATCCCAGAAATAAATCTGGAGTATAAGGGCAAGG<br>CCCCGAAGAAAATCAAGCTGAACATAGAAGCACCGGAGGGAGCGCATCTGACGA<br>TTGGCGAGATCAACATAACGGCCGAAAATTCCGATGTAGAGGAAATTGAAGTGAAT<br>GTAAAAGGCAACGTAACCATAAACAAACTTACGATCAAAGCGATTAACTCAAAGATT<br>AAGAAAATCAACGTTAAACTTGAGGGGAAAGTGACCGTACCTGAATTAGAAATTAT<br>TGCGGAGAACAGCGAAGTCGAGGAAATCAATGTGGAGGTGGAAGGGGATGTGA<br>CTGTCCCTAAAATTACAATTAAAGCTATCAATAGTAAGATAAAGAAGATCACGGTTAA<br>GCTCGAGGGTAAAGTTACCGTGCCAGAGATAGAGATTATAGCCGAGAACTCGGAA<br>GTTGAGGAGATTAATATCGAATTGGAAGGTGATGTTACAGTACCCAAGATCAACCTT<br>ATCGCGGAAAACTCTACCATTAAGAAATTTACTCTGAAGATTGAAGGTAATGTTACT<br>GTTGGCAAAATCATAGTGGAGAATAAGGAAGGATCTAAGCCTATAGATGTCGAAATT<br>ATCATCAAGAAGAACGGTGAAAACGTCACCAAGAAGTACACTGTGAACGAAGGC<br>GAAACCCTGATCATTGAGAACCTCATTAACAAGCCGtaa |
| **Bcap_2_2** | ATGATGGGCAGCAGCCACCATCACCACCACCACTCCAGCGGTCTTGTGCCTCG<br>GGGTAGCagtGACGAAGAGGAGCTGACAAAGAAACTAGAAGAGGGCGAAGTCGA<br>GATAAAAGGCAAATACGGGGAGATAAAAGTCAAAGGAAAGGTAGAGCTCAAAGAT<br>GGAACCGCCAAAGTGGTGTTTACGGATGATGGCAAGGTGTCGTTAGAGATATCAG<br>GCAATGCGATCATTGAGGATAGCGAGATCGAAGTAAAGGGTAAGAATGGGAAGGA<br>GGTCGGCATCGAATTCAGCGGCAACATAACCATAAAGAACTCGAAAATTAAAGTTA<br>CTGGAGACAACATAGAAAAGGCCTATATTAAAATCAGTGGGAACGCCACGATCGA<br>GAACAGTGAAATAGAGGTGAAGGGGACCAACTCGGAGGAAGTATATATTGAAATAA<br>GCGGTAACGCGACAATCAAGGACTCTGAGATTAAGGTCACCGGAACTAACTCCA<br>AGAAGGTTTACATCAAAATTTCTGGAAACGCTACAATTGAAAACAGCGAAATCGAG<br>GTTAACGGCACCAATTCTGAAGAAGTTTATATCGAAATATCCGGGAATGCTACTATC<br>AAGAATAGCACAATTAGTGTGACGGGCACGAACAGCAAGAAAGTGTACATTAAGAT<br>TGAGAACATTTCCGAAGCAGGGACGGCGACTATTACGATAACCGAGACTGGTAAA<br>ACTGAAACCAAGGATGTCAAGGCTGGTGATTCTCTGTATTATGAGATCAACAACCC<br>TAAGAACGGGAAAATAACTGTGAACTTCAAAGCTAAGGCGtaa |
| **Bcap_3_1** | ATGATGGGCAGCAGCCACCATCACCACCACCACTCCAGCGGTCTTGTGCCTCG<br>GGGTAGCagtGCGGCGATCACCCCAGAAGCGAAAGCGGCCATAGAGAAGTTAAA<br>AGCCGCGGGTGCCAAACTCACGATTCCAGACACGGTGGACGGACTCAAGATTA<br>GCAAAGTGGAGGTAAAAGACGGGAAGGTGACTATAGAAGAGATTACAGCTACTTC<br>CAAAACCACGGCCGGGATCGAGATCTCAAACCTGAAAGCGGAGGAAATAAACAT<br>CAATAAGATAAAAGTAACCGCAGGTGTAACGGCATCTGTTACCCTGTCGAACATAA<br>AGGCGAAGAAAATAAATATAAAGAAAATTGAAGTGACAGCAGGCGTGACTGCAAG<br>CATTAAGTTGAGCAATATAGAGGCGGATGAGATAAACATAGATGAAATTAAGGTCAC<br>AGCCGGCGTAACTGCGAGCGTCACTCTGTCTAACATTAAAGCTAAGAAGATTAACA<br>TCAACAAAATCGAGGTCACCGCCGGTGTTACCGCGTCTATTAAACTGTCCAACAT<br>CGAAGCTGACGAAATCAACATCAAGGAGATCAAGGTGACCGCTGGCGTTACTGC<br>CAGTATTGACTTTAGTAATATCAAAGCAGACAAGTTCAACATTGACAAGATTGAGGT<br>TACCATAACCAAGGGTGGTACGGCTAAAATCAGTGGCAACTTCGAGGGCAAGGC<br>CGAAGGCAAATTTAACATCACTAAGGAACTGAAGCCTGGTGAAACCCTCATTGTG<br>AAGATCAAAGATGGGAAACTGGAGTCTGTAGAACTAATTAAGCCGtaa |

| Supplementary Table 3: DNA sequences of designed β-solenoid solenoid proteins with caps that were experimentally characterised | |
|---|---|
| **Design name** | **DNA sequence** |
| **Bcap_3_2** | ATGATGGGCAGCAGCCACCATCACCACCACCACTCCAGCGGTCTTGTGCCTCG<br>GGGTAGCagtATGAACACCCTGGAGGAGCTCAAGGAACACTTAGAAAGCCTCGGT<br>TTTAAAGTCAAAATAGAGGGCGATACCTTGTATTTCGAATACGAGGACGAAAACGG<br>CGGGAAACTGAAAGGCAAAGTAAAAGTTGAGAATGGCAAGATCACGGTGGAGTT<br>TGAGGCGGATTCGTTAGAGAAGTTTACGCTGTTTGAAAATTTCACGTTCAACGCAA<br>AGAGCGTGGAGATAAAGGGCAAATTGGACAGCCCAAAAGAACTTACGATTTTCAA<br>GAATAGCACATTCAATGCAGACTCAGTCACCTTTGAGTTCGAAGCGAAGTCCCCT<br>AAAGAGATGACGTTATTTAAGAACAGTACATTTAACGCTAATAGTGTGACGATCAAG<br>GGTAAATTAGATAGCCCGGAGAAGCTGACAATCTTCGAGAACTCTACGTTTAACGC<br>CAAATCTATCAAGCTAGAGCTGGAAATGGATAATCCGAAAGAAATGACCATCTTTAA<br>GAATTCCACAATTAATGCCGATAACGTTGACATAAAACTGTCTATAAAGAACCCAGA<br>AAAGCTTACACTCTTCAAGAACACGACCCTCAATGTTAAGTCGCTGAATATTACCAT<br>TACTGATAAGGAGACCGGCGAAACCTGGGAAGGGACCTTCACTGGTACGGGCA<br>CTCTGAACATCACCATAACTAAGAAGGATGGCGAGTATACTATCGAGGTAGAAACA<br>ACGGGTGGTATAACCGGTGAGATTAAGAAGGTTGAAtaa |
| **Bcap_4_1** | ATGATGGGCAGCAGCCACCATCACCACCACCACTCCAGCGGTCTTGTGCCTCG<br>GGGTAGCagtATGAACACATTGGAGGAACTAAAAGAGCATCTGGAATCGCTGGGG<br>TTTAAAGTTAAAATTGAGGGCGACACCCTGTATTTCGAATACGAAGACGAAAACGG<br>CGGGAAGCTGAAAGGCAAGGTCAAAGTGGAGAATGGCAAGATCACTGTGGAGTT<br>TGAAGCCGATAGCCTAGAAAAGTTCACCTTATTCGAAAACTTCACATTTAACGCTAA<br>ATCGGTGGAAATAAAGGGCAAATTAGACAGCCCTAAGGAGCTGACCATTTTCAAG<br>AATAGCACGTTTAATGCAGATAGTGTTACCTTTGAGTTCGAAGCGAAATCTCCGAA<br>AGAAATGACCCTATTTAAGAACAGCACCTTCAATGCCAACAGCGTGACGATCAAG<br>GGGAAACTAGATTCGCCGGAAAAGCTCACCATATTTGAGAACTCGACATTCAACG<br>CCAAATCCATTAAGCTTGAGCTAGAGTTGGACTCGCCTAAAGAGATGACAATCTTC<br>AAGAATTCTACTATAAACGCGGATTCGGTAGATATCAACATTAGCATAAAGAACCCA<br>AAGAAACTGATTCTGTTTAAGAACACCACGTTCAACGTCAATACCCTTAACATCAA<br>AATCACGGACAAGGAGACTGGTAAAACCTGGGAAGGGACTTTCACGGGCACCG<br>GCACTCTGAACATAACCATCGAGAAGAAGGACGGGGAGTACACGATTACCGTCT<br>CTACGACCGGAGGAATTACTGGCAGTATTACGGAAGTCAAAtaa |
| **Bcap_4_2** | ATGATGGGCAGCAGCCACCATCACCACCACCACTCCAGCGGTCTTGTGCCTCG<br>GGGTAGCagtATGATGAAGACGGAGGAATTATTAGCCAAACTGAAAGCAGCGGGC<br>CTGACGGGCAAGGTGGAGTACGATGAAGCGACCAATAGCATTACCGTGGAAGCC<br>TCGGGTACTGTGGCTGAGGGGAAGAAAGTGACGATAGAGCTGCCGGCCGATGT<br>GAAGAAGGCTACTCTGACCCTAAAGAACGTGACCATTAAGAAGAACGCGAAACTA<br>ACCGCAGAGGTAAAGGCGACAGATACCGCGACCTTAAACGTAGAAAACTTGACG<br>CTAGAAGAGAATGCGGAAGTGAACTTACGTGTAGAGGCCAAGACTACAGCAACC<br>CTGCGACTCAAGAATGTGACTCTCAAAGATAACGCCAAGCTTACTGCAGAAGTCA<br>AATCCGAAAAGACTGCCACCCTGGAGGTTGAAAATCTAACGTTGGAAAACAACAG<br>CGAAGTAACTTTGCGCGTGAAAAGCGGAGACACAGCTACGATTAACCTGAAGAAT<br>GTTACACTTAAGAATAACTCTAAGCTGAATATTGAAATCGAATCTGGTAAGACCGCT<br>ACCTTGAACTCGAAGAACCTTCGTCTCGAGGAGGGCGTTGAGGTCAACATAACA<br>ATTACGCTGAACGGCAAAACCATCACAATCACTATTACTCGCGATTCTAATGGCAAT<br>CTCCATATGAGTATTACAGTCGACGCGGGAGTGGATACACTGACTATAGATGTAAA<br>AGAGGAAAACGGATATTTGAATTTAACGATCACCGTTACGGCGGCGtaa |

| Supplementary Table 3: DNA sequences of designed β-solenoid solenoid proteins with caps that were experimentally characterised | |
|---|---|
| **Design name** | **DNA sequence** |
| **Bcap_5_1** | ATGATGGGCAGCAGCCACCATCACCACCACCACTCCAGCGGTCTTGTGCCTCG<br>GGGTAGCagtATGCTGACGGCAGCCGAGATAGAAAAGAAAGCCAAAGAGCTGGG<br>CTATAATGTCACGATTACAGGTACGAGCACAGGCGGCAAAGTGCCTACCGGGACT<br>GTGAGCGAAAAGGATGGCAACGTAACGATCACCATAAATCTGCCCTCGGGCGCC<br>TCCAGCTTTACTGGAAACATAACAGTGTCCAATGTGAAAGGAAAGAAAATCACTAA<br>CAATGTCAATATAAACGTTGGGAAATTCACGGGCAATATCACCTTTAGCAATATAAC<br>GGCGGAAGAGATCGAGAACAACGTAAACATTAATGCGAAGAAGTTCGAAGGGAAT<br>ATTACCTTCTCAAATATAACTGCGGATAAAATTAATAATAACGTCACAATCAACGTCG<br>ATGAGTTCATTGGTAACATCACGTTCAGCAACATTACCGCTAAGGAGATTAACTTGT<br>CTGTCGAAATAAACGCGAAGAAATTTAAGGGGAACATCAACTTTTCCAACATAACC<br>GCGGATGAAATCAACATAAATGTAAATATTAACGTGGAGAAGTTTGAAGGCAAGATT<br>ATCTTTAGTAACATTACGGCTAAGAAGATCAACCTGAATATTACTTTCAAGGGCCTC<br>AAGGACGGGGAGAAGCTTAAACTGACCCTAACTAATGAGAAAACGGGTGAGACT<br>CTGACTAAGACGGAGACCGCCAAGAACGGTGAAGCGACCATTAACATCAATGGG<br>GCGAATGGCTACAGCGGGGTGGCCGAAGTAGAGAAAtaa |
| **Bcap_5_2** | ATGATGGGCAGCAGCCACCATCACCACCACCACTCCAGCGGTCTTGTGCCTCG<br>GGGTAGCagtAGCACGACCATAACCAAAACCATTGAATTAGGCGATAAGAATGAAG<br>TGAATATCAACAACATTTCGCTGACGGGCGACAACGTCACCCTGATTATTAAAGGT<br>AGTAAAGAGAAAATAAAGAAGGTCACCATCAACTCTATAAACGTGACTAGCACAAC<br>CACGGCGAAAGTTATCATCGAAAACATCAAGGCAGACAAAATAAATATTAATGAAAT<br>AAATGTGACGGCCGGGAAGACCGCTAAAATAGAAATTAAGAATATCGAAGCCAAG<br>GAGATAAACATTAACAAAATTAACGTAACAGCGGGAGAGACCGCAGAAATTAAAAT<br>TGAAAACATAAAAGCGGATAAGATTAACATCAATGAGATCAATGTCACGGCTGGTAA<br>AACTGCGAAGATCGAGCTGAGCAACATTAAGGCCAAAGAAATCAATATAAACAAAA<br>TCAACGTGAAAGCAGGGGAAACCGCCGAAATTAAAATCAAGAACATAGAGGCGG<br>ACAAGATCAATATCAACGAGATTAATGTAACCGCGGGTAAGACTGCTGTGGTGGAA<br>TTCGAGAATATCAAAGCTAAGGAAATTAACATTAACAAGATCAACATTACGATTGTG<br>GATGCCGACGGTAACGAGGCTACGTATACTGGCAAAGCGACCGGAAAGAATGGC<br>AAGCTGGAGAAGATAGAGATGGAAAAGTCGAGTGGCAGCCTGAACACGGAACTG<br>AAGATTGAGAACGGCAAAATCACCGCGACTCTGAAAtaa |
| **Bcap_6_1** | ATGATGGGCAGCAGCCACCATCACCACCACCACTCCAGCGGTCTTGTGCCTCG<br>GGGTAGCagtGCCCTGCAGCTTACTCTGGACCTGACTTACAAGGAGAACCCGGG<br>CGAAGGCGGAGGTAAACTTACGTTGGAGGATGGGGCCGGAAACAAGACCACTAT<br>AAACTTTCCAGCCAATGTGTCTAAAATCACCATCAAAGAGATTAGTGTGACGGGTG<br>CCACGACGGCGTCCGTAACCCTGTCCAATATAACCGTGGACGAGTTGAATATCGA<br>AAAGATAGAGGTGACAGCGGGAAATACGGCCAGCATTGTCCTCAGCAACATAAAG<br>GCTGACAAAATAAATATCAAGGAAATCAAAGTAACGGCAGCGAATACTGCGAGTAT<br>AGAGCTATCCAACATCACTGCTAAAGAGATCACGATTAAGAAAATAGAAGTGACCG<br>CGGGGAACACCGCTAGCATCAAACTGAGCAATATCAAAGCGGATAAGATCAACAT<br>CGAGGAGATAAAAGTGACTGCGGGTAACACAGCGTCGATTGAGATTTCTAACATA<br>GAGGCAGACGAAATCAATATTAAGAAGATCGAAGTTACAGCCGGCAACACGGCTA<br>CTATCAAGTTGTCGAACATTAAAGCTAAGAAAATTAACATCGAGAAAATTACGATAAA<br>TGCCGCGAACGGTACAATTATTCTTGAAAACATTGAGGGCAAAGGCAAGATTACCA<br>TTGAAGATGGCGGCGAGACGTTAACTATTGAAATTAATGGGAAGATTGAAAAGAAG<br>AAAATTGAAATTAAGGGCGGTAAGATTACACTGATTGAAtaa |

| Supplementary Table 3: DNA sequences of designed β-solenoid solenoid proteins with caps that were experimentally characterised | |
|---|---|
| **Design name** | **DNA sequence** |
| **Bcap_6_2** | ATGATGGGCAGCAGCCACCATCACCACCACCACTCCAGCGGTCTTGTGCCTCG<br>GGGTAGCagtGCGGCGAAAGGCTCGATCACAATTAGTGGGGAAGTCAAGGGAGG<br>CAGCGCAACCGTGAAAGTAGAACTGAAAGACTCCAAGGGTAAGGTGACGAAGTA<br>TGAGAAGGAGTACAAGGACGGAGAGACTTTCGAGTTGACAATCGAAGGGGCGGA<br>TGACATTACCGAAATAAATATAGAGATCAAGGCAACGGGTGGTTTTACCCCGAACA<br>TAACCCTAAAGAACATAAAGAATTTAAAGAAGATAAACATCAAAATAGAAGCAGACA<br>AAGATCTGACGCCCAACTTGACCATTGAAAACGTGGAAGGTCTTAAGGAAATCAA<br>CATTAACATGAAGGTTAAGAAGAACCTCAAACCTAATATAACGATGAAGAACGTCAA<br>AGGCCTGAAGAAGATTAACATCAACATGGAGGTCGACGAGGACCTGACACCAAAT<br>ATTACATTGGAAAATGTCGAGGGACTGGAAGAGATTAATATCAACATGAAGGTGGG<br>CAAGAATCTAAATTTGAACATCAATATGAAGAATGTGAAAGGGCTCAAGAAAATTAA<br>CATCAATATCACCGTAGGCGAGACGCTGAATATTAACATTACGTATAAGAACGTAGA<br>TGATTTTAAGTTAAACATCACGATTAAAGCGACCAAGCCGCTAAAAGGTACCATCA<br>CTATTAAGAATGGCGATCCTAACAAAGAAACAATAACAGAGCCAATTGAGTTAGAG<br>GAGGGTGAGACACTCACTCTGAACTACGATATAAAAtaa |
| **Bcap_7_1** | ATGATGGGCAGCAGCCACCATCACCACCACCACTCCAGCGGTCTTGTGCCTCG<br>GGGTAGCagtATCAAGATAACGGTGGAGGGCGAGGGTGTTTCGGGCACCATAAAG<br>ATCTATGATGAGAATGGCAACCTGCTGTACGAGAAGAAATTTGAGACCAAGCCAG<br>GCGAAAAGAAGAAAACCGTGGAAATTGAGCTGCCGGAAGACGCCAAAGACATCA<br>GCAAGATCGAAATCAATATCAGTTCTAAGACTGGCCCACTCAAAGTGAATATAAACA<br>TAGAAAACTTGAATGTCAAGGAGATCAATATCAACGTGTCCTCGGAAAAGCAGCC<br>ACTGACTGTCGATATCAACATTAAGAACGTCAACGCGGATAAAATAAACATCAACAT<br>CTACTCTGAGGAAGGTCCGCTTACTGTGAACATTAATATAGAGAACGTAAATGTGA<br>AAGAGATAAACATAAACATCAAGAACAAGAAAGCTCCCCTCACCGGCGAGATTAC<br>CATTGAAAATGTAAAAGCCGATAAGATTAACATTAATATTTATAGCGAAGAGGCCCC<br>GCTGACAATAAACATAAATATTAAGAATGTCAATGTGGACAAAATTAACATCAACGTT<br>ACTAATAAGAAAGCGCCCGCGAACGTTAACATAACCGCGGAGAACGTGAATTCCA<br>AGGAAACGACCATCACTATTACAGACAAGGGCAAAACAACTACGGTCAAAATCGA<br>AGGCGGCGGGAACTTTAAAATTACGCAGAAGGACGGTACCTTAAACGTCGAGACT<br>AAGGAAGGGGTTAAGAAAGTTGAAGTAAAGGAGGAAtaa |
| **Bcap_8_1** | ATGATGGGCAGCAGCCACCATCACCACCACCACTCCAGCGGTCTTGTGCCTCG<br>GGGTAGCagtATGGAACTGACCGCCGAAGAGTTCATCCAGAAGGCAAAAGCCGG<br>GGAAACCATAGAAAACGCGACGGTGGATAACGTAGAGATAAAGAATGAAAAGTTTA<br>GCCAAGTGACCTTTAAGAACGTGGATTTCAAGAACGTTAAGTTTGAGTCGGTAAA<br>GTTCGAAAACGTGCGCTTCGAAAATGTCCGGTTCGACAACGTGAGATTTGAAACT<br>GTACGGTTTGAGAATGTAAAATTCCGCAACGTAAAATTCGAGAACGTTAAATTCAG<br>AAATGTCAAGTTTGAAAACGTACGCTTTGAGAGTGTGGACTTTAAACAGGTTAAAT<br>TTGACAATGTTAAATTCGATAATGTGCGATTTAGATCAGTGAAATTTGATAACGTCAA<br>ATTTCGTAACGTCCGTTTTGAATCTGTTCGTTTCGACAATGTGAAATTTGAGACTGT<br>CACCTTCGAATCGGTGAAGTTCGAGTCTGTGAAAGCAGATAAGGGTGTAGAAGTG<br>AAGAACGTGAAGGGCCTTACCAAAGAGGAGCTGGAGAAGATATTTAAGtaa |
| **Bcap_8_2** | ATGATGGGCAGCAGCCACCATCACCACCACCACTCCAGCGGTCTTGTGCCTCG<br>GGGTAGCagtATGGAGCTGACCGCCGAAGAGTTTATCCAGAAAGCAAAAGCCGG<br>CGAGACCATTGAGAACGCCACAGTAGAAAACGTAGAGATTAAGAACGATCGCTTC<br>AGCAGCGTCACCTTCCGCAACGTGGATTTTAAGAACGTCAAGTTCGAGAATGTGA<br>AATTTGAAAACGTGCGCTTCGAAAATGTAAAATTTGACCAAGTCCGGTTTGAAAAT<br>GTGCGATTCGAAAACGTCCGCTTTAAATCTGTTCGTTTTGAACAAGTTAGATTCCG<br>TAATGTGAAATTCGATACGGTGAAATTCGAAAATGTGGAGTTCAAGCAGGTCCGAT<br>TTGAAAATGTACGCTTTGATTCAGTGAAATTCAAGAACGTGAAGTTTGATCAGGTTA<br>AATTTAAGAACGTTAAATTTGATAACGTAAAGTTCGAAAATGTTAAGTTTGAAAACGT<br>TACCTTTGAGCAAGTGCGTTTCGACACGGTAAAGGCCGACAAAGGGGTGGAAGT<br>AAAGAATGTCAAGGGACTGACGAAGGAGGAACTGGAGAAGATATTTAAAtaa |

| Supplementary Table 4: Data processing parameters for Bcap_8_2 crystal |  |  |  |
|---|---|---|---|
| | **Overall** | **Low** | **High** |
| **High resolution limit** | 10.05 | 27.17 | 10.05 |
| **Low resolution limit** | 117.74 | 117.75 | 10.22 |
| **Completeness** | 100 | 100 | 100 |
| **Multiplicity** | 5.7 | 4.2 | 6.2 |
| **I/sigma** | 3.6 | 4.7 | 0.9 |
| **Rmerge(I)** | 0.266 | 0.167 | 1.098 |
| **Rmerge(I+/-)** | 0.247 | 0.157 | 0.968 |
| **Rmeas(I)** | 0.292 | 0.193 | 1.2 |
| **Rmeas(I+/-)** | 0.292 | 0.191 | 1.149 |
| **Rpim(I)** | 0.118 | 0.093 | 0.478 |
| **Rpim(I+/-)** | 0.156 | 0.108 | 0.614 |
| **CC half** | 0.981 | 0.987 | 0.761 |
| **Wilson B factor** | 727.47 | | |
| **dF/F** | 0.153 | | |
| **dI/s(dI)** | 0.444 | | |
| **Total observations** | 7768 | 390 | 433 |
| **Total unique** | 1355 | 93 | 70 |
